# The kinesin KIF1C transports APC-dependent mRNAs to cell protrusions

**DOI:** 10.1101/2020.11.30.403394

**Authors:** Xavier Pichon, Konstadinos Moissoglu, Emeline Coleno, Tianhong Wang, Arthur Imbert, Marion Peter, Racha Chouaib, Thomas Walter, Florian Mueller, Kazem Zibara, Edouard Bertrand, Stavroula Mili

## Abstract

RNA localization and local translation are important for numerous cellular functions. In mammals, a class of mRNAs localize to cytoplasmic protrusions in an APC-dependent manner, with roles during cell migration. Here, we investigated this localization mechanism. We found that the KIF1C motor interacts with APC-dependent mRNAs and is required for their localization. Live cell imaging revealed rapid, active transport of single mRNAs over long distances that requires both microtubules and KIF1C. Two color imaging directly showed single mRNAs transported by single KIF1C motors, with the 3’UTR being sufficient to trigger KIF1C-dependent RNA transport and localization. Moreover, KIF1C remained associated with peripheral, multimeric RNA clusters and was required for their formation. These results reveal an RNA transport pathway in mammalian cells, in which the KIF1C motor has a dual role both in transporting RNAs and in promoting their clustering within cytoplasmic protrusions. Interestingly, KIF1C also transports its own mRNA suggesting a possible feedback loop acting at the level of mRNA transport.

## Introduction

Localization of mRNA to specific subcellular compartments is an important mechanism for the spatio-temporal regulation of gene expression in diverse cell types and organisms (Chin and Lécuyer, 2017; Eliscovich and Singer, 2017). Subcellular mRNA localization allows localized protein synthesis, which is important for many biological functions such as cell fate determination (Berleth et al., 1988), cell polarization (Condeelis and Singer, 2005), cell division (Chouaib et al., 2020), cell migration (Katz et al., 2012; Wang et al., 2017; Moissoglu et al., 2020), embryonic patterning (Forrest and Gavis, 2003) and synaptic plasticity (Martin and Zukin, 2006; Lin and Holt, 2007). One of the best characterized examples is the yeast *Ash*1 mRNA that localizes specifically in the bud of the daughter cells and encodes a transcriptional repressor protein involved in suppressing mating-type switching (Paquin and Chartrand, 2008). Studies of this and other models revealed that the subcellular localization of mRNA relies on three main mechanisms, acting separately or in combination: random diffusion combined with local entrapment, general transcript degradation coupled to localized protection and directed transport along the cytoskeleton (Cody et al., 2013; Medioni et al., 2012; Bovaird et al., 2018).

Diffusion of the mRNA to a localized anchor is used for example by *nanos* mRNA, which localizes to the posterior pole of the *Drosophila* egg during late oogenesis (Zaessinger et al., 2006; Jeske et al., 2011). This localization mechanism is based on facilitated diffusion, promoted by the cytoplasmic streaming generated by motor proteins in the oocyte (Ganguly et al., 2012), and trapping, achieved by an actin-dependent association with the germ plasm at the posterior pole (Forrest and Gavis, 2003; Kugler and Lasko, 2009). A non-uniform distribution of transcript can also be obtained by a targeted degradation mechanism. In *Drosophila*, *hsp83* mRNA is uniformly distributed in the fertilized egg, but as nuclear divisions progress, total RNA levels decrease drastically by global degradation, and only the *hsp83* mRNAs localized at the pole plasm are protected from degradation (Semotok et al., 2005, 2008). This differential stability of *hsp83* mRNA is regulated by distinct cis-acting elements. Hsp83 degradation element (HDE) and Hsp83 instability element (HIE) are jointly involved in the degradation pathway by recruiting the RNA binding protein (RBP) Smaug, which then binds the CCR4/POP2/NOT deadenylase complex to set off the degradation of *hsp83* mRNAs (Semotok et al., 2005). Hsp83 protection element (HPE), present in the 3’UTR downstream of the HDE, is sufficient to protect *hsp83* transcript in the pole plasm (Semotok et al., 2005). Finally, active, motor driven transport of mRNAs along the cytoskeleton is a very common localization mechanism. This mechanism generally involves cis-acting elements, also called zipcodes, contained in the 3’UTR sequence of the transcript. This is exemplified by the case of the β-actin mRNA in vertebrates, which accumulates at the leading edge of migrating cells and was among the first localized mRNAs discovered (Singer, 1993). This mRNA contains a zipcode sequence recognized by the RNA Binding Protein ZBP1, allowing the transport of β-actin mRNAs in a motor-driven manner along the cytoskeleton (Kislauskis et al., 1993; Oleynikov and Singer, 2003; Liao et al., 2015; Condeelis and Singer, 2005). Interestingly, transport of β-actin mRNA by ZBP1 involves both microtubules (MTs) and actin filaments (Fusco et al., 2003; Oleynikov and Singer, 2003), and several motors appear involved in its transport, with some cell type and compartment specificity. Indeed, MYO5A and KIF5A interact with ZBP1 to transport β-actin mRNAs in dendrites and axons (Ma et al., 2011; Nalavadi et al., 2012), while Myosin IIB (MYH10) and KIF11, which directly binds ZBP1, regulate the transport of β-actin mRNAs in fibroblasts and during cell migration (Song et al., 2015; Latham et al., 2001). Interestingly, it has been shown that ZBP1 orthologues in different organisms are able to bind mRNAs other than β-actin, such as Vg1 mRNA that localizes at the vegetal pole of Xenopus oocytes (Havin et al., 1998). These ZBP1 orthologs are involved in regulating mRNA localization, translation or stability (Yisraeli, 2005; Jønson et al., 2007; Vikesaa et al., 2006).

Apart from β-actin, numerous other RNAs are localized at protrusions of mesenchymal cells and their local translation is important for cell migration (Mili et al., 2008; Mardakheh et al., 2015; Moissoglu et al., 2020; Costa et al., 2020). Localization of these RNAs is carried out through at least two distinct pathways. First, a subset of about a hundred RNAs, which include transcripts encoding signaling and cytoskeleton regulators (such as the Rab GTPase RAB13, the RhoA exchange factor NET1, the collagen receptor DDR2, the motor related proteins TRAK2, DYNLL2, and others), require the APC tumor suppressor protein for localization and have been referred to as APC-dependent (Wang et al., 2017). Second, protrusion-enriched RNAs, exemplified by RNAs encoding ribosomal proteins, do not require APC and exhibit distinct regulation (Wang et al., 2017).

Similar to what has been described for other localized RNAs, sequences within the 3’UTR of APC-dependent RNAs are necessary and sufficient for targeting to the periphery (Mili et al., 2008). Specifically, interfering with or deleting particular GA-rich regions is sufficient to disrupt peripheral localization and perturb cell movement in various systems (Moissoglu et al., 2020; Costa et al., 2020; Chrisafis et al., 2020). Furthermore, localization to the periphery requires the microtubule cytoskeleton and in particular a subset of stable, detyrosinated microtubules(Wang et al., 2017; Moissoglu et al., 2019). Indeed, at least some APC-dependent RNAs exhibit a co-localization with the plus ends of detyrosinated microtubules (Mili et al., 2008). The peripheral complexes also contain APC, a protein that has the ability to directly bind microtubules via its C-terminus (Barth et al., 2008; Bahmanyar et al., 2009; Jimbo et al., 2002; Munemitsu et al., 1994; Zumbrunn et al., 2001), hence suggesting that APC might mediate the interaction of localized mRNAs with microtubules (Mili et al., 2008; Preitner et al., 2014).

An additional feature integrated with the localization of APC-dependent RNAs is their existence in distinct physical states. In particular, RNAs in internal or peripheral, actively extending cytoplasmic regions exist as single molecules that are undergoing translation. However, at some peripheral areas single RNAs coalesce in multimeric heterogeneous clusters that are composed of multiple distinct RNA species. Interestingly, these clusters preferentially form at retracting protrusions and contain translationally silent mRNAs (Moissoglu et al., 2019). These data indicate the existence of a dynamic regulatory mechanism during cell migration, which coordinates local mRNA translation with protrusion formation and retraction. However, the exact mechanisms and molecular players involved in transport to the periphery and cluster formation for this group of RNAs are still unclear.

In this study, we focused on the KIF1C mRNA and protein, which we recently showed accumulate and colocalize in cytoplasmic protrusions (Chouaib et al., 2020). We describe a specific mRNA transport mechanism by which the KIF1C kinesin motor binds APC-dependent mRNAs, including its own, actively transports them to cell protrusions in a 3'-UTR dependent manner and additionally participates in promoting and/or maintaining their peripheral clusters.

## Results

### Identification of a specific mRNA subset localizing with KIF1C motor in human cells

High-throughput mRNA-protein cross-linking approaches previously showed that KIF1C directly binds mRNAs (Baltz et al., 2012; Castello et al., 2012), and we recently showed that KIF1C mRNAs and proteins colocalized together in protrusions of HeLa cells (Chouaib et al., 2020) Figure 1A), suggesting that the KIF1C kinesin might be somehow involved in the metabolism of protrusion mRNAs. To determine the identity of the mRNAs bound by the KIF1C motor, we used a HeLa cell line stably expressing a KIF1C-GFP fusion from a bacterial artificial chromosome containing all the regulatory sequence of the human KIF1C gene, including its 5’ and 3’ UTRs (Poser et al., 2008; Chouaib et al., 2020). We immunoprecipitated (IP) KIF1C-GFP with anti-GFP antibodies or uncoated beads, as controls, and identified the co-precipitated RNAs using microarrays (Figure 1B and Table S1). We found that many mRNAs were enriched in the KIF1C-GFP IP as compared to the control IP. A GO term analysis of the top 200 mRNAs associated with the KIF1C-GFP motor revealed an enrichment for “post-Golgi vesicle-mediated transport” (5.4 fold enriched, pV 3 10^−2^), “organelle localization by membrane tethering” (4.2 fold enriched, pV 8.10^−3^), “microtubule-based process” (3.6 fold, pV 9 10^−7^), and “cilium assembly” (3.5 fold, pV 1.3 10^−4^). To explore in more detail the localization of the mRNAs associated with KIF1C, we performed a small smFISH localization screen in HeLa cells using 26 of the mRNAs most enriched in the KIF1C IP (Table S1 and S2). These included the RAB13 mRNA (5.7 fold enrichment, Table S1), along with the KIF1C mRNA itself (2.6 fold enrichment) and the NET1 and TRAK2 mRNAs (5.7 and 4.9 fold enrichment respectively), which were reported to localize to protrusions in mouse cells in an APC-dependent manner (Wang et al., 2017). Visual examination of the images revealed that KIF1C, NET1, TRAK2 and RAB13 mRNAs clearly localized to protrusions of HeLa cells (Figure S1A). Quantification of the mean mRNA distance to the cellular membrane confirmed their localization, which was not observed for three controls mRNAs (KIF20B, PAK2, MYO18A; Figure S1D). To confirm the link between APC-dependent mRNA localization and binding to KIF1C protein, we performed a correlation analysis of the two metrics (Figure S1E). This indicated that APC-dependent mRNAs indeed preferentially associate with KIF1C protein, while mRNAs coding for ribosomal proteins, which often localize to protrusion independently of APC (Wang et al., 2017), do not. The IP/microarray data thus show a physical link between KIF1C and mRNAs that localize to protrusions in an APC-dependent manner.

**Figure 1:**
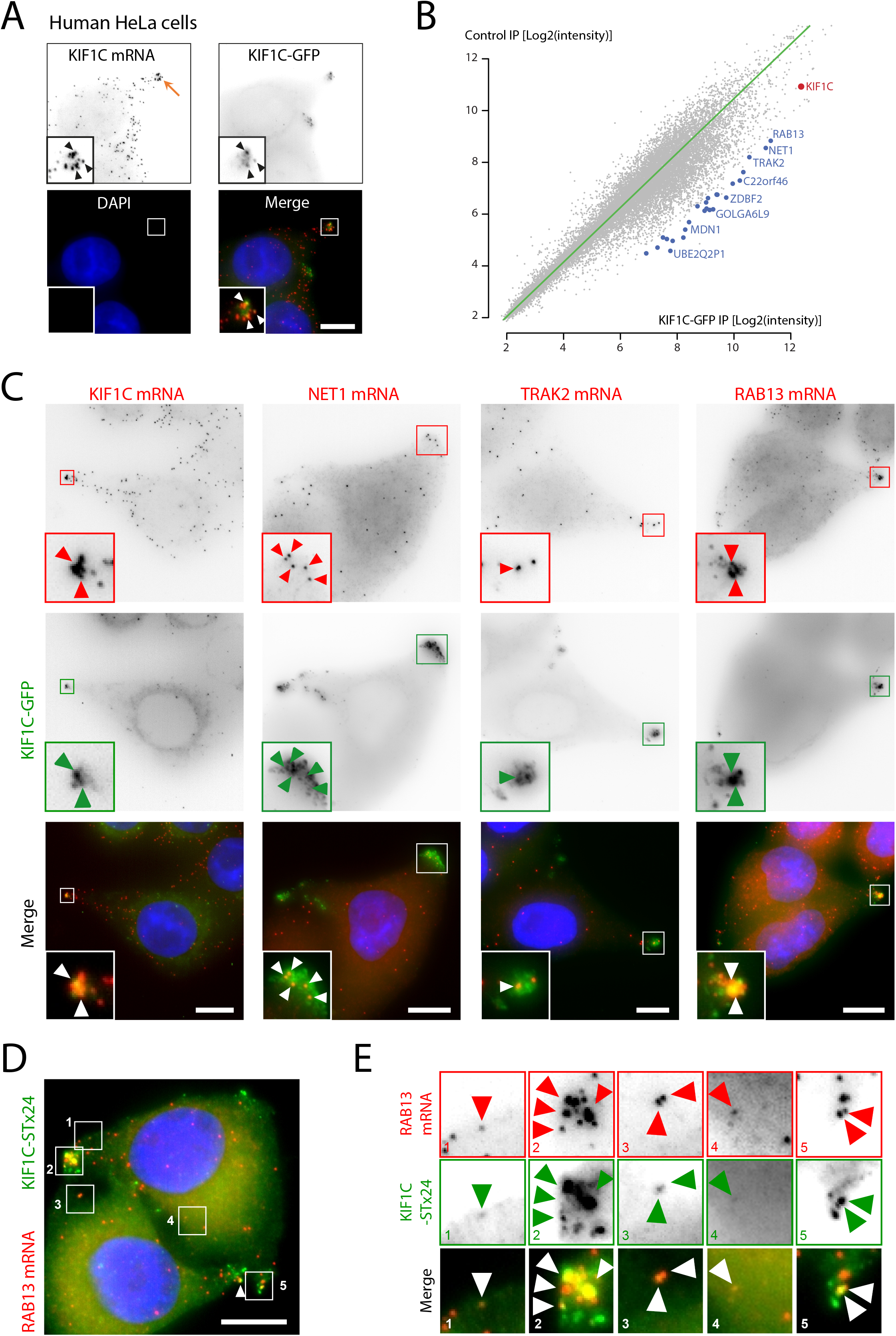
Identification of mRNAs associated with the KIF1C motor. **A-** The KIF1C kinesin colocalizes with its mRNA in protrusions of HeLa cells. Images are micrographs of a H9 FlipIn HeLa cell line stably expressing a KIF1C-GFP cDNA. Top left: KIF1C mRNA detected by smFISH with probes against the endogenous mRNA; top right: KIF1C-GFP protein; bottom left: DNA stained with DAPI, bottom right: merge of the two signals with the KIF1C-GFP protein in green and KIF1C mRNA in red. Orange arrow: a cell protrusion. Blue: DNA stained with DAPI. Scale bar: 10 microns. Insets represent zooms of the boxed areas in the merge panel. White and black arrowheads indicate the colocalization of KIF1C mRNA with KIF1C-GFP protein. **B-** Transcripts associating with the KIF1C-GFP protein. The graph depicts the microarray signal intensity of RNAs detected in a KIF1C-GFP pull-down (x-axis), versus the control IP (y-axis). Each dot represents an mRNA. Red dot: KIF1C mRNA; blue dot: mRNAs enriched in the KIF1C-GFP IP. **C-** Colocalization of KIF1C-GFP with KIF1C, NET1, TRAK2 and RAB13 mRNAs. Images are micrographs of HeLa H9 Flip-In cells stably expressing a KIF1C-GFP cDNA, labelled by smiFISH with probes against the indicated mRNAs. Top: Cy3 fluorescent signals corresponding to endogenous KIF1C, NET1, TRAK2 and RAB13 mRNAs. Middle: KIF1C-GFP signal. Lower: merge with the Cy3 signal in red and GFP signal in green. Blue: DNA stained with DAPI. Scale bar: 10 microns. Arrowheads indicate accumulation of single mRNA molecules at cell protrusions. **D-** Single molecule colocalization of KIF1C-ST_x24_ with RAB13 mRNAs. Images are micrographs of Hela cells stably expressing a KIF1C-ST_x24_ expression and scFv-GFP. Red: Cy3 fluorescent signals corresponding to RAB13 mRNAs labeled by smiFISH with probes against endogenous RAB13 mRNA. Green: GFP signal corresponding to single molecules of KIF1C protein. Blue: DNA stained with DAPI. Scale bar: 10 microns. **E-** Insets represents zooms of the numbered areas from panel D. Legend as in D. Arrowheads indicate molecules of RAB13 mRNA and KIF1C-ST_x24_ protein.

Next, we tested whether these mRNAs colocalize with the KIF1C protein *in vivo*. To this end, we performed smFISH experiments in a HeLa cell line stably expressing a KIF1C-GFP mRNA from a cDNA and found that indeed, KIF1C, NET1, TRAK2 and RAB13 mRNAs co-localized with the KIF1C-GFP protein in cytoplasmic protrusions (Figure 1C). In order to show that this colocalization reflected a molecular interaction, we performed single molecule imaging using the SunTag system (Tanenbaum et al., 2014). To this end, we generated a stable cell line expressing KIF1C-fused to 24 repeats of the GCN4 epitope (KIF1C-ST_x24_), together with the single-chain variable fragment fused to sfGFP (scFv-sfGFP). This system enables the detection of single molecules of the KIF1C protein in live cells (Tanenbaum et al., 2014; Figure 1D), and we combined detection of single KIF1C-ST_x24_ proteins with single mRNA detection by smFISH using probes against RAB13 mRNA (Figure 1D and 1E). KIF1C-ST_x24_ protein and RAB13 mRNAs were found to colocalize at protrusions as expected (Figure 1E, panels 2-5). In addition, we also observed colocalization events at the single molecule level at more internal locations in the cytoplasm (Figure 1E, panels 1, 3-4). This confirmed the interaction of single molecules of KIF1C protein with single molecules of RAB13 mRNAs. Taken together, these data suggest that the kinesin KIF1C might be part of a mechanism that localizes APC-dependent mRNAs to cytoplasmic protrusions.

### KIF1C is required for the localization of APC-dependent mRNAs to cytoplasmic protrusions in human and mouse cells

To test whether the localization of mRNAs in protrusions depended on the KIF1C protein, we depleted KIF1C by siRNA in HeLa cells, and performed smFISH experiments using probes against RAB13 mRNA. Remarkably, RAB13 mRNAs became less localized when KIF1C expression was reduced with siRNAs (Figure S1B-D), demonstrating that the KIF1C kinesin was required for RAB13 mRNA localization in human cells.

Next, we moved to a mouse system, NIH/3T3 cells, where the localization of mRNAs in protrusions has been extensively studied (Chicurel et al., 1998; Kislauskis et al., 1997; Mingle et al., 2005; Mili et al., 2008; Wang et al., 2017; Moissoglu et al., 2019). To test whether the KIF1C protein has a general role in localizing mRNAs at cell protrusions, we assessed the localization of a series of APC-dependent and APC-independent mRNAs by smFISH, following depletion of KIF1C expression with two different siRNAs. Intracellular distributions of mRNAs were quantitatively assessed by calculating a Peripheral Distribution Index (PDI), a metric that distinguishes diffusely distributed from peripherally localized RNAs (Wang et al., 2017; Stueland et al., 2019). A PDI value of 1 indicates a uniform, diffuse signal, while values smaller or greater than 1 indicate a perinuclear or peripheral localization, respectively. As shown in Figure 2A-B, several APC-dependent RNAs, including *Net1, Rab13, Ddr2, DynII2* and *Cyb5r3*, exhibited a protrusion localization pattern that was lost following KIF1C depletion (Figure 2A and Figure S2A). This change in mRNA distribution was reflected by a lower PDI value, characteristic of a uniform or more perinuclear distribution (Figure 2B and Figure S2B-C). Interestingly, the localization of two APC-independent mRNAs, *Rps20* and *Rpl27a*, was not affected (Figure S2C). To ascertain that this effect was not due to altered mRNA expression, we measured their levels following KIF1C depletion (Figure S2D). This analysis showed no changes in the overall abundance of APC-dependent mRNAs, except for the depleted KIF1C mRNA (Figure S2D). Therefore, we conclude that KIF1C is required for the localization of APC-dependent mRNAs to cell protrusions, in both human HeLa cells and mouse NIH/3T3 fibroblasts.

**Figure 2:**
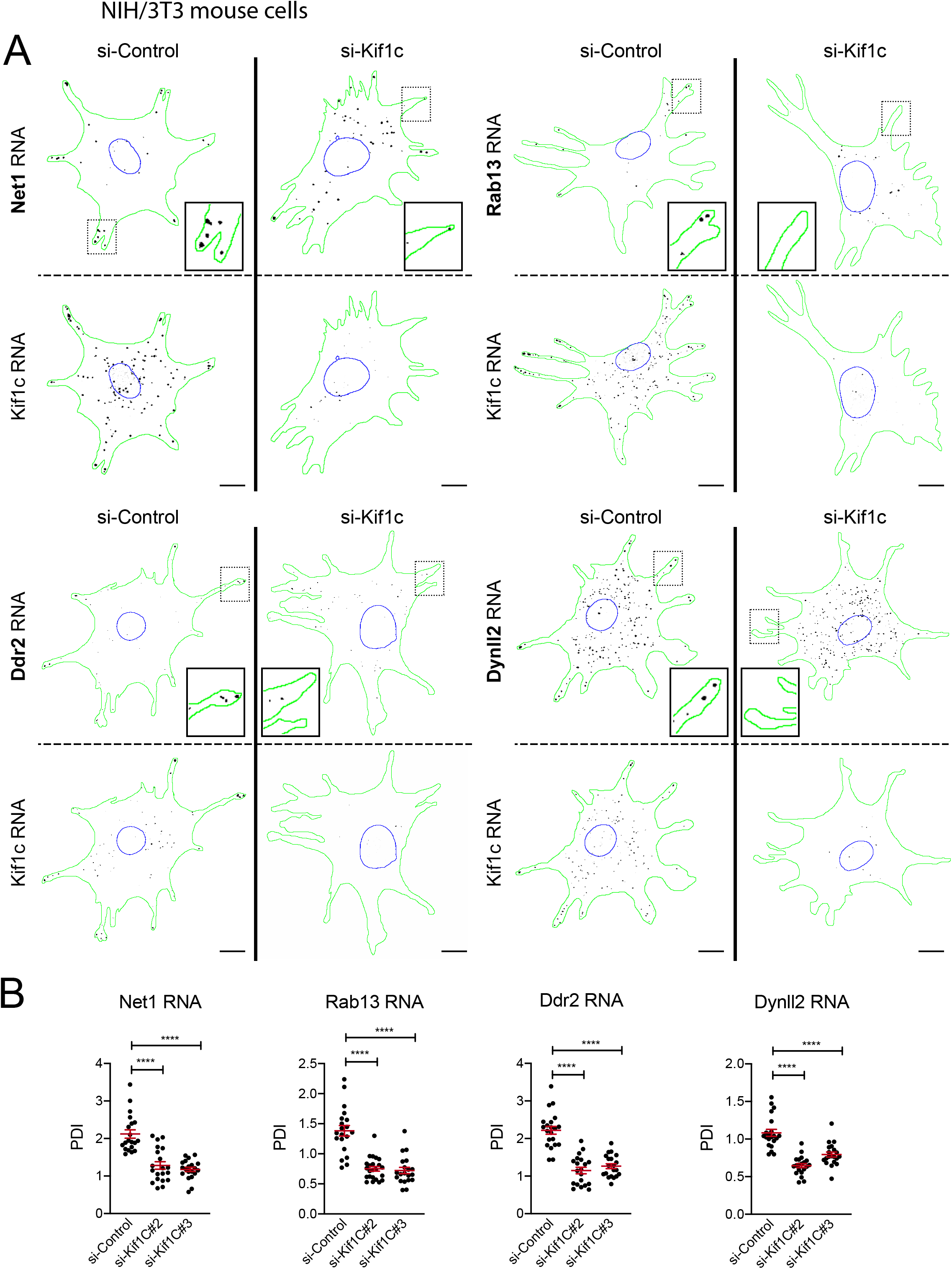
KIF1C is required for the localization of APC-dependent mRNAs to cytoplasmic protrusions in mouse fibroblasts. **A-** Depletion of Kif1c prevents mRNA accumulation in protrusions. Images are micrographs of NIH/3T3 cells labelled by smFISH with probes against *Net1*, *Rab13*, *Ddr2*, *Dynll2* and *Kif1c* mRNAs, following treatment with siRNAs against Kif1C (panels si-Kif1c) or a control sequence (panel si-Control). Scale bars: 10 microns. Green: outline of the cells; blue: outline of the nuclei; black: smFISH signals. Insets represent magnifications of the boxed areas. **B-** Quantification of mRNA localization of cells described in A. Graphs represent the intracellular distribution of the indicated mRNAs as measured by PDI index, with and without treatment of cells with the indicated siRNAs. Red bars represent the mean and 95% confidence interval. Points indicate individual cells. Stars are p-values: ****<0.0001, ***<0.001, estimated by analysis of variance with Bonferroni’s multiple comparisons test.

### KIF1C is required for active transport of APC-dependent mRNAs on microtubules

To monitor trafficking of APC-dependent mRNAs, we expressed in NIH/3T3 fibroblasts a reporter carrying the β-globin coding sequence followed by 24 binding sites for the bacteriophage MS2 coat protein (MCP; Figure 3A). Binding of co-expressed MCP-GFP to these sites allows visualization and tracking of single molecules of the reporter mRNA in living cells (Fusco et al., 2003). To recapitulate the localization of APC-dependent RNAs, the reporter additionally included a control 3’ UTR or the 3’ UTR of *Net1* or *Rab13* (hereafter referred to as β24bs/Ctrl, β24bs/Net1 and β24bs/Rab13, respectively). As shown previously, these 3’ UTR sequences are sufficient to direct peripheral distribution of this reporter transcript in NIH/3T3 cells (Moissoglu et al., 2019). We initially examined trafficking of the reporter during early stages of cell spreading, which mimic conditions in actively protruding cell regions. Indeed, live fluorescence imaging of the reporter containing the Net1 3’ UTR revealed a distinct peripheral pattern after plating cells on fibronectin for 30 minutes (Figure 3B and Movie 1). Because kinesin-dependent mRNA trafficking is expected to occur on the microtubule cytoskeleton, it was important to identify microtubule-dependent events and discriminate them from other modes of motion. For this, reporter particles were tracked in cells before and after 15min of nocodazole treatment, which depolymerizes microtubules (Figure 3B and Movies 1, 2). To identify long and linear movements, as those expected to occur on microtubules, we used two different metrics, ‘Linearity of forward progression’ and ‘Track displacement’ to quantitatively describe individual tracks. This analysis revealed a subset of tracks (3-6% of the total tracks) that exhibits higher values for these parameters in control cells (net displacement > 4 microns, linearity > 0.7), and which were absent in cells treated with nocodazole (Figure 3C, D, F and Movies 1, 2). In contrast, a control reporter lacking a localizing 3’ UTR did not produce tracks with these characteristics (Figure 3E, F). Importantly, this subset of tracks was not affected by disruption of the actin cytoskeleton with cytochalasin D or following treatment with a control vehicle, DMSO (Figure 3C, D and Movies 3, 4). Tracking of a reporter carrying the Rab13 3’ UTR also exhibited long linear tracks with dependence on microtubules (Figure 3E, F). Thus, the reporters carrying the 3’ UTR of *Net1* or *Rab13* mimic the localization pattern of APC-dependent mRNAs and allow the identification of long and linear microtubule- and 3’ UTR-dependent transport events.

**Figure 3:**
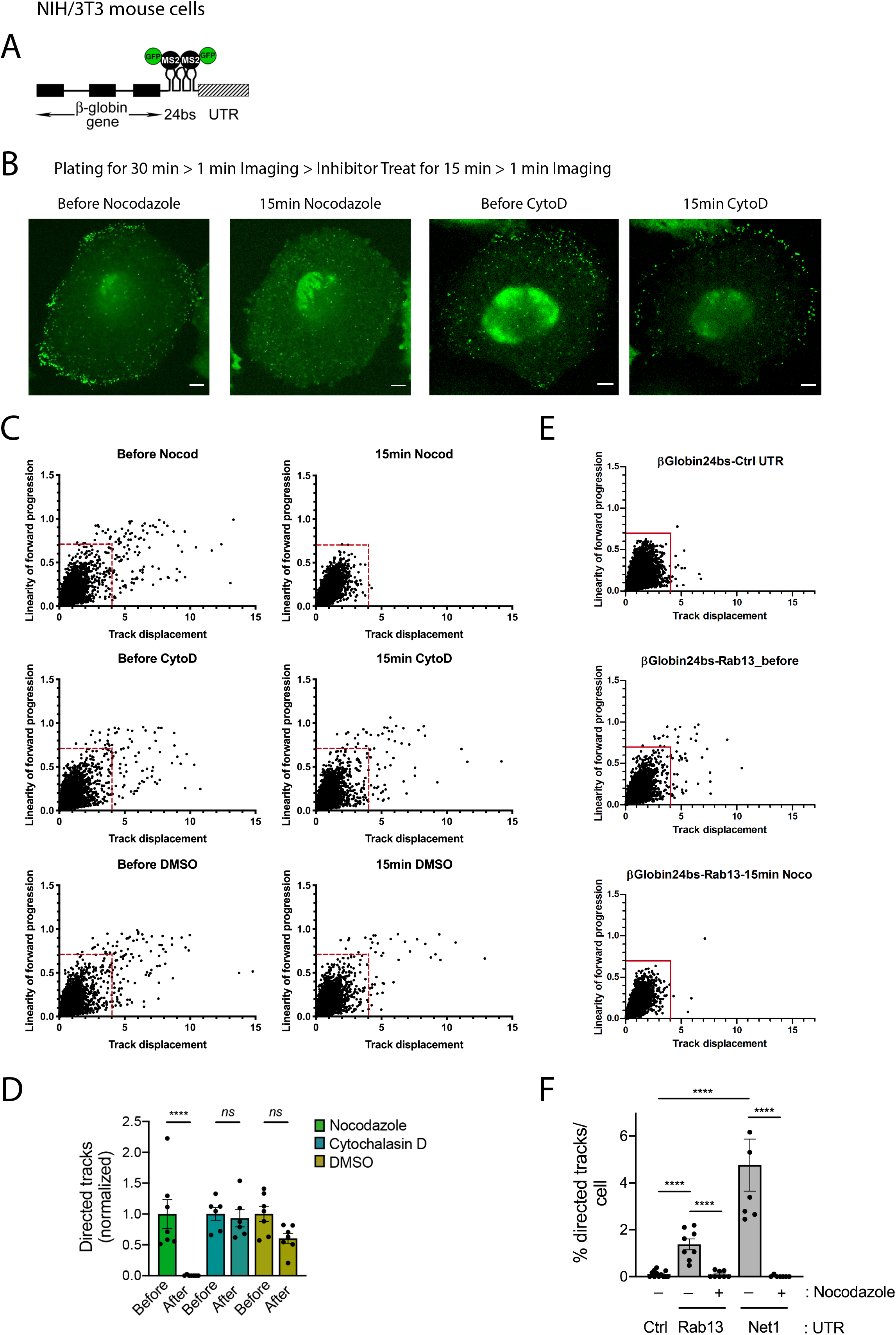
Reporter mRNAs containing Net1 and Rab13 3’UTRs display long, linear microtubule-dependent displacements. **A-** Schematic of the mRNA reporter construct containing the β-globin coding sequence followed by 24xMS2binding sites and the indicated 3’UTRs (β24bs/3’ UTR). **B-** Images are snapshots of live NIH/3T3 cells taken 30 minutes after plating. High speed imaging was performed over 1min to track individual RNA movements before and after15 minutes of drug treatment with Nocodazole or Cytochalasin D as indicated. See movies 1-4 for time lapse imaging. The cells stably expressed the β24bs/Net1 reporter mRNA and MCP-GFP (Green). Scale bar: 5 microns. **C-** Single RNA molecules of the β24bs/Net1 reporter mRNA were tracked over a 1min period in cells treated as described in B. The graphs plot the displacements of individual tracks (x-axis) over the linearity of their forward progression (y-axis), (defined as the mean straight line speed divided by the mean speed). Red lines indicate the thresholds used to filter tracks of molecules undergoing directed movement. N=6-7 cells. **D-** The bar plot depicts the percentage of directed tracks per cell (β24bs/Net1 reporter mRNA) before and after treatment with the indicated compounds. Average values of respective ‘Before’ values were set to 1. Stars represent p-values: ****<0.0001, ns: non-significant, estimated using one-way analysis of variance with Tukey’s multiple comparisons test. Error bars: standard error of the mean. **E-** Single RNA molecules of the β24bs/Control UTR (Ctrl) or the β24bs/Rab13 UTR reporters were tracked over 1min period in cells treated or not with nocodazole. Graphs plot the displacements of individual tracks (x-axis) over the linearity of their forward progression (y-axis). Red lines indicate the thresholds used to filter tracks of molecules undergoing directed movement. N=8-12 cells. **F-** The graph depicts the percentage of directed tracks of the indicated reporters per cell following treatment with nocodazole. Stars represent p-values: ****<0.0001, estimated using one-way analysis of variance with Sidak’s multiple comparisons test. Error bars: standard error of the mean.

To directly test the role of KIF1C in these trafficking events, we visualized fluorescent particles of the reporter carrying the *Net1* 3’ UTR and measured the frequency of long and linear, microtubule-dependent displacements in actively spreading cells following KIF1C depletion. As previously observed with endogenous transcripts (Figure 2), reporter mRNAs became less localized when KIF1C expression was reduced with siRNAs (Figure 4A and Movies 5-8). Importantly, track analysis showed that KIF1C loss significantly reduced the number of the microtubule-dependent displacements (Figure 4B and 4C). To assess specificity, we depleted two additional kinesins that have been linked to APC and RNA transport, KIF5B and KIF3A (Dunn et al., 2008; Kanai et al., 2004; Cai et al., 2009; Yasuda et al., 2017; Baumann et al., 2020). Figure 4A-C shows that depleting these kinesins did not change the overall peripheral accumulation of the reporter and did not result in reduction of the long and linear transport events. Thus, KIF1C exhibits a specific function in transporting APC-dependent mRNAs via microtubules in actively protruding cell regions.

**Figure 4:**
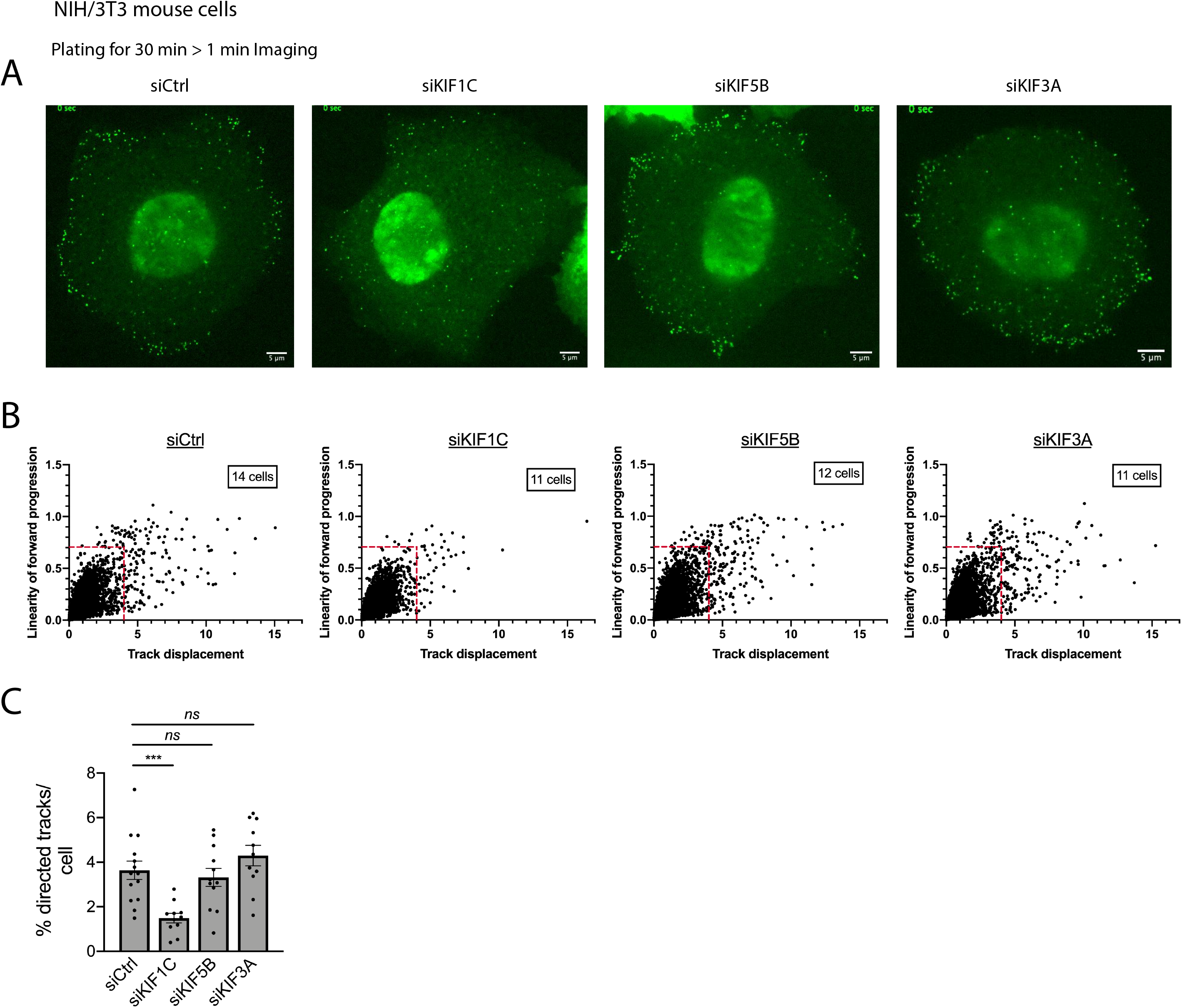
Reporter mRNAs containing the Net1 3’UTR require Kif1c for long, linear microtubule-dependent displacements. **A-** Images are snapshots of live NIH/3T3 cells taken 30 minutes after plating. The cells stably expressed the β24bs/Net1 mRNA reporter and MCP-GFP (Green) and were treated with the indicated siRNAs. The green spots correspond to single mRNAs detected with the MCP-GFP. High speed imaging was performed over 1min to track individual RNA movements. See movies 5-8 for time lapse imaging. Scale bars are 5 microns. **B-** Graphs plot the displacements of individual RNA tracks (x-axis) over the linearity of their forward progression (y-axis), (defined as the mean straight line speed divided by the mean speed), using the movies of cells as shown in A. Red lines indicate the thresholds used to filter tracks of molecules undergoing directed movement (based on Figure 3). **C-** Graph depicts the percentage of directed tracks per cell following treatment with the indicated siRNAs (see panel B). Stars represent p-values: ***<0.001, ns: non-significant, estimated using one-way analysis of variance with Dunnett’s multiple comparisons test. Error bars: standard error of the mean.

### Peripheral clustering of APC-dependent mRNAs depends on KIF1C

Peripheral APC-dependent mRNAs can form large heterogeneous clusters that are translationally silent (Moissoglu et al., 2019). These clusters often associate with retracting protrusions in migrating cells, suggesting that they are part of a spatio-temporal control of protein synthesis (Moissoglu et al., 2019). Formation of these clusters is recapitulated by the reporter constructs carrying the *Net1* or *Rab13* 3’ UTR, but not by a control reporter (Figure S3). These clusters are visible at later time points after plating (ca. 3 hours), when most protrusions are not actively extending, consistent with the appearance of endogenous RNA clusters in non-extending or retracting protrusions (Figure 5A; Moissoglu et al., 2019). These clusters can be identified as bright particles with intensities higher than those characteristic of single molecules (Figure 5A). To test whether KIF1C is implicated in the formation of these clusters, we scored the frequency of bright particles in KIF1C-depleted cells during late stages of spreading (3 hours; particle brightness > 4950). As shown in Figure 5A, while clusters formed by the *Net1* 3’ UTR-reporter were readily observed in protrusions of control siRNA-treated cells, their frequency was substantially reduced, and mostly single molecules were present, when KIF1C was depleted (Figure 5A-C). Moreover, cluster formation was only marginally affected by the depletion of KIF5B or KIF3A. Thus, KIF1C specifically controls the clustering of APC-dependent mRNAs. We note that clusters are not detected even in protrusions containing a substantial amount of single RNA molecules (see enlarged KIF1C insets in Figure 5A), suggesting that cluster loss is not a secondary consequence of reduced number of mRNA molecules arriving at protrusions upon KIF1C depletion. We rather think that these results indicate an additional role of KIF1C in forming higher order RNP complexes at protrusions.

**Figure 5:**
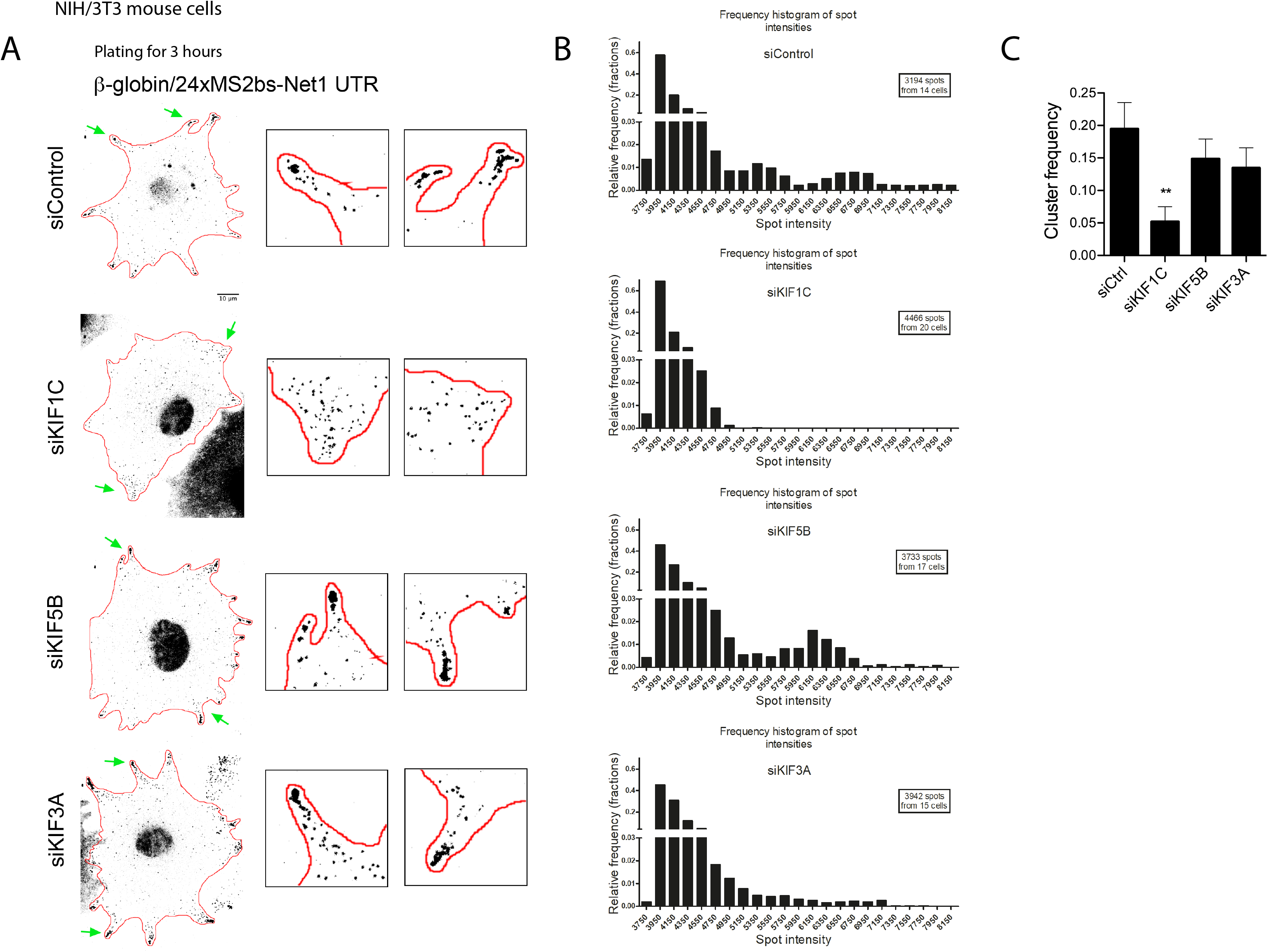
Kif1c is required for the peripheral clustering of reporter mRNAs containing the Net1 3’UTR in live mouse fibroblasts. **A-** Images are micrographs of live NIH/3T3 cells taken 3h after plating and expressing MCP-GFP and the β24bs/Net1 reporter mRNA. Cells were treated with the siRNAs indicated on the left. Scale bar is 10 microns. Boxed insets are magnifications of the areas indicated by green arrows. **B-** Frequency histograms of the intensities of the β24bs/Net1 reporter mRNA spots following treatment with the indicated siRNAs, measured from images as shown in A from N=14-20 cells. **C-** Graph depicts the mRNA cluster frequency following treatment of cells with the indicated siRNAs. Clusters correspond to β24bs/Net1 mRNA spots of intensities higher than 4,950, measured from the graphs shown in B. Stars represent p-values: **<0.01, estimated using one-way analysis of variance with Dunnett’s multiple comparisons test. Error bars: standard error of the mean.

### Single molecule two color imaging provides direct evidence that the KIFIC1 motor transports protrusion mRNAs

To provide direct evidence that the protrusion mRNAs are transported by the KIF1C motor, we performed two color single molecule imaging of mRNAs and motors, in order to visualize co-transport of the two types of molecules. To this end, we used the NIH/3T3 cells expressing the Net1 3’ UTR-containing reporter and modified them to also stably express a KIF1C protein fused to the SunTag (KIF1C-ST_x24_), together with a single-chain variable fragment antibody fused to mScarletI (scFv-mScarletI).

Imaging of fixed cells showed that the KIF1C-ST_x24_ motor and reporter mRNAs accumulated in protrusions as expected (Figure 6A-B, Figure S4A-B). Moreover, we could also occasionally detect colocalization events where a single molecule of KIF1C-ST_x24_ would colocalize with a single reporter mRNA at internal cellular areas. To confirm that this colocalization was relevant to mRNA transport, we performed two-color live-cell imaging using movies recorded at a high frame rate (7.36 fps for 52 seconds). This allowed the detection of co-transport events, in which a single molecule of KIF1C-ST_x24_ moved with a reporter mRNA molecule in a rectilinear manner at high speed (Figure 6C, D; Movie 9; Figure S4C, D; Movie 10). Kymographs confirmed that both molecules moved together in an anterograde direction toward protrusion (Figure 6E and Figure S4E), traveling an average distance of 22 microns at speeds of 2.6 μm/second (Figure 6F-G). Taken together, these data demonstrate that the KIF1C kinesin actively transports this *Net1* reporter mRNA to cell protrusions along microtubule cables.

**Figure 6:**
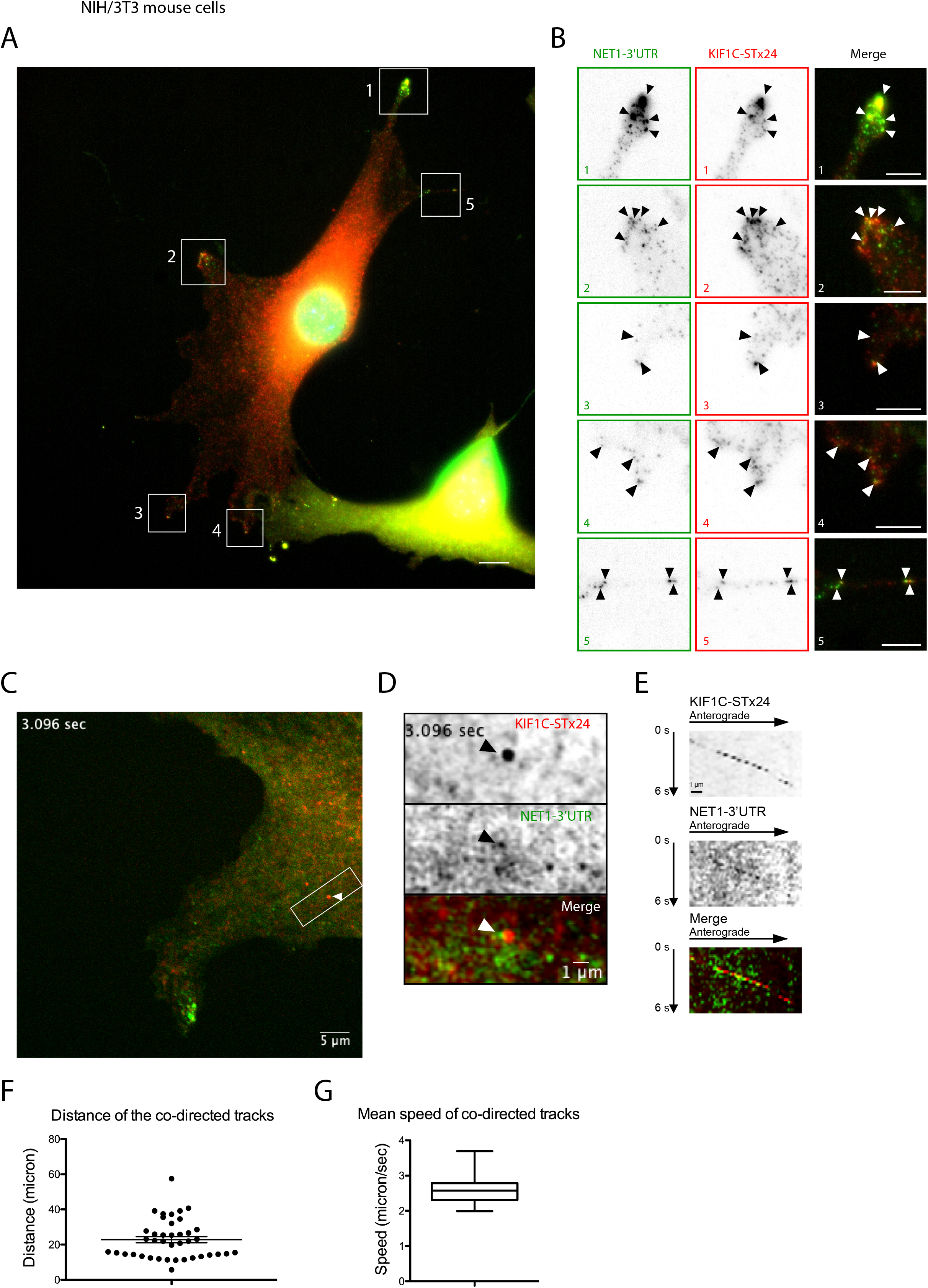
The KIF1C motor transports mRNAs containing the Net1 3’UTR to cell protrusions. **A-** Images are micrographs of fixed NIH/3T3 cells expressing the β24bs/Net1 reporter mRNA, MCP-GFP (green), KIF1C-ST_x24_ protein and scFv-mScarletI (red). Single molecules of the β24bs/Net1 reporter mRNA are visible in green, while single molecules of KIF1C-ST_x24_ protein are red. The numbered white boxes are magnified in B. Blue: DNA stained with DAPI. Scale bar is 5 microns. **B-** Insets represent magnifications of the boxed areas from the image shown in A. Left: MCP-GFP signals labelling β24bs/Net1 mRNAs; middle: scFv-mScarletI labelling KIF1C-ST_x24_ protein; right: merge with mRNAs in green and KIF1C-ST_x24_ in red. Black and white arrowheads indicate colocalization of single molecules of β24bs/Net1 mRNAs and KIF1C-ST_x24_. Scale bar is 5 microns. **C-** Snapshot of a live NIH/3T3 cell expressing β24bs/Net1 mRNA, MCP-GFP (green), KIF1C-ST_x24_ protein and scFv-mScarletI (red). Snapshot is extracted from Movie 9. The white arrowhead indicates a co-transport event of a single molecule of β24bs/Net1 mRNA (green) with a KIF1C-ST_x24_ protein (red). The boxed area is magnified in panels D and E. Scale bar is 5 microns. **D-** Magnification of the boxed area in panel C, highlighting a co-transport event. Top: KIF1C-ST_x24_; Middle: β24bs/Net1 mRNA; bottom: merged panel with the β24bs/Net1 mRNA in green and KIF1C-ST_x24_ in red. Scale bar is 1 micron. **E-** Kymograph from Movie 9, showing the trajectory of a single molecule of KIF1C-ST_x24_ (top panel), a single molecule of β24bs/Net1 mRNA (middle panel) and the merge (bottom panel). The cellular area shown corresponds to panel D. **F-** The graph depicts the distance travelled by co-transported molecules of KIF1C-ST_x24_ and β24bs/Net1 mRNA. Each dot is a track, and the mean and 95% confidence intervals are shown by horizontal lines. **G-** Boxplot depicting the mean speed of co-transported molecules of KIF1C-ST_x24_ and β24bs/Net1 mRNA in NIH/3T3 cells. Speed is microns/second. The vertical bars display the first and last quartile, the box corresponds to the second and third quartiles, and the horizontal line to the mean.

## Discussion

RNA transport along the cytoskeleton is a well-established mechanism allowing subcellular mRNA localization and local translation. In mammals, a large class of mRNAs localize to cytoplasmic protrusions of many cell types, where they are anchored at the plus-end of detyrosinated microtubules by APC. Here, we show that these mRNAs associate with the microtubule motor KIF1C and interestingly, they include the KIF1C mRNA itself. We show that APC-dependent mRNAs and KIF1C protein co-localize in protrusions and can also be seen co-transported together along directed tracks. Moreover, the peripheral localization of these mRNAs as well as their microtubule-dependent motion depend on KIF1C, demonstrating that it is an essential motor that transports APC-dependent mRNAs to protrusions. Our data provide a striking *in vivo* visualization of the co-transport of individual RNA molecules with a specific molecular motor, involved in a widespread RNA transport pathway.

### The kinesin KIF1C controls a widespread mammalian mRNA transport system

RNA localization controls spatial and temporal aspects of gene expression in a variety of species and cell types. Although its significance is better understood in specialized cells such as neurons, recent reports highlight its widespread prevalence, including in cells with a mesenchymal phenotype (Mardakheh et al., 2015; Wang et al., 2017; Fazal et al., 2019; Costa et al., 2020; Chouaib et al., 2020). Our current understanding of the transport mechanisms includes the requirement of *cis*-acting sequence elements and *trans*-acting factors, which work with the actin or microtubule networks and motor proteins to bring mRNAs to their destination (Cody et al., 2013; Medioni et al., 2012; Bovaird et al., 2018). Nevertheless, our knowledge of common RNA transport mechanisms that operate in most, if not all cells, is limited. In the case of protrusion mRNAs, which localize in all cell types examined so far, their localization was shown to require APC and detyrosinated microtubules (Mili et al., 2008; Wang et al., 2017). It is not yet known whether APC is transported together with the KIF1C-RNA complexes or whether it is independently transported and subsequently associated with RNAs at the periphery. *In vitro* studies suggest that motile complexes can be formed between mRNA, APC and KIF3A (Baumann et al., 2020). Our data, however, clearly show the involvement of KIF1C in transporting protrusion mRNAs *in vivo*. Moreover, APC was shown to accumulate at the leading edge of migrating cells using kinesin-1 and kinesin-2 (Mimori-Kiyosue et al., 2000; Nakamura et al., 2001; Ruane et al., 2016). The use of distinct motors suggests an independent transport for APC and protrusion mRNAs. It is also possible that this diversity of motors reflects differences between cell types, as has been described for the β-actin RNA that uses different motors in neurons and fibroblasts (see Introduction; Ma et al., 2011; Nalavadi et al., 2012; Song et al., 2015; Latham et al., 2001). One potential reason for using different motors could pertain to the dynamics and transport speeds required in each case. In this context, the fact that KIF1C appears to be the fastest human cargo transporter (Lipka et al., 2016) might provide an advantage that could underlie its preferential use in the highly dynamic mesenchymal cells. Furthermore, kinesins other than KIF1C, such as KIF5B, contribute to localization of APC-dependent RNAs (Yasuda et al., 2017). We speculate that they may function in specialized cells or affect different aspects of localization that are distinct from transport *per se*. In line with this idea, protrusion mRNAs display a complex translational regulation concomitant to the protrusion dynamics and a local reorganization of RNA clusters (Moissoglu et al., 2019), likely requiring a number of yet uncharacterized actors.

Another important question deals with the adaptors that link KIF1C to mRNAs. On one hand, proteomic data showed that KIF1C protein interacts with the exon junction complex (EJC; Hein et al., 2015). The EJC is assembled on spliced RNAs and serves as an interaction platform for proteins that direct mRNA export, localization, translation and nonsense-mediated mRNA decay (NMD), suggesting a role for the EJC in the transport of protrusion mRNAs. On the other hand, regions rich in G and A nucleotides are present in the 3’UTR of APC-dependent RNAs and are part of the localization element (Moissoglu et al., 2020; Costa et al., 2019). Furthermore, the 3’UTR is sufficient to direct protrusion localization of intronless exogenous constructs (Moissoglu et al., 2019, 2020), indicating that a potential KIF1C recruitment through exon-exon junctions might not be necessary for protrusion localization. Specific RNA-binding proteins could serve as adaptors that mediate KIF1C recruitment similar to the model suggested for other RNA transport complexes. However, given that high-throughput mRNA-protein cross-linking approaches previously showed that KIF1C directly binds mRNAs (Baltz et al., 2012; Castello et al., 2012), an alternative interesting possibility would be that in this case the motor directly selects and binds to its RNA cargo.

### KIF1C triggers mRNA clustering in cytoplasmic protrusions

Our live imaging experiments show that reporter mRNAs are transported predominantly as single molecules during the early stages of cell spreading, reminiscing the single molecule appearance of RAB13 RNA in actively protruding regions in migrating cells (Moissoglu et al., 2019). Remarkably, APC-dependent mRNAs coalesce into higher order clusters at peripheral regions during later time points of cell spreading and this phenotype depends specifically on KIF1C. We think that clustering is not merely a consequence of peripheral mRNA accumulation, because we have not observed it in the absence of KIF1C even at protrusions occasionally containing substantial numbers of single mRNA molecules. We rather favor the explanation that clustering is a separate function of KIF1C that is temporally and spatially regulated. Given that these clusters contain stably anchored mRNAs (Mili et al., 2008), we envision that KIF1C switches from a microtubule motor to an mRNA anchoring module promoting clustering. A similar switch has been observed in Drosophila oocytes, whereby Dynein converts from a motor of *gurken* mRNA to a static anchor at its final destination (Delanoue et al., 2007). Such a switch on KIF1C may take place on pre-existing motor molecules as they reach the periphery or may be a function of newly-synthesized KIF1C translated from its peripherally localized mRNA. It is still unclear how such a switch would occur and/or whether it might additionally involve a change in the RNA-binding mode of KIF1C (direct or indirect through other RNA-binding proteins). Clusters of APC-dependent mRNAs have been previously reported to be heterogeneous and to contain translationally silent mRNAs (Moissoglu et al., 2019). Thus, overall our results point to a spatially and temporally controlled mRNA clustering role of KIF1C that is separate from its motor function and that might be coordinated with translational regulation.

### RNA transported by KIF1C mediates diverse functions at cell protrusions

Peripheral localization of APC-dependent RNAs promotes cell migration (Wang et al., 2017). Specifically, approaches targeting the localization elements of these mRNAs, as a group (Wang et al., 2017) or individually (Moissoglu et al., 2020), resulted in inhibition of cell migration. These effects are likely due to a requirement for locally translating these mRNAs for full activation of the encoded proteins (Moissoglu et al., 2020). Interestingly, KIF1C has been shown to control adhesion dynamics and cell migration (Theisen et al., 2012). It was proposed to act via the trafficking of α5β1 integrins, and our results indicate that the transport of APC-dependent mRNAs to the periphery likely contributes to the mechanism by which this kinesin controls cell migration. Along this line, the GO terms associated with the top 200 KIF1C-associated mRNAs presented in this study (i.e. post-Golgi vesicle-mediated transport; organelle localization by membrane tethering; microtubule-based process; cilium assembly) indicate how KIF1C-mediated mRNA transport could impact processes related to cell motility.

### KIF1C protein localizes its own mRNAs to cell protrusions: a transport feedback loop

The KIF1C transcript localizes to cytoplasmic protrusions in mammalian cells. Moreover, it co-localizes with KIF1C protein in protrusions (Chouaib et al., 2020), and we show here that the KIF1C protein physically associates with its own mRNA. This local accumulation of KIF1C could be involved in an RNA clustering and anchoring mechanism as discussed above, but it could also serve to transport additional mRNAs by alternating back-and-forth movements on the cytoskeleton. Indeed, locally translated KIF1C protein would allow the motor to explore the local cytoplasm and transport back additional mRNAs to protrusions using the same MT tracks. Such a bidirectional motility has been reported for KIF1C and it is mediated by the scaffold protein Hook3. This protein forms a complex between dynein and KIF1C (Kendrick et al., 2019), and regulates their activities to allow the motor to perform multiple transport cycles while avoiding a tug-of-war between opposite motors (Siddiqui et al., 2019). The fact that KIF1C also brings its own mRNA to protrusions suggests the possible existence of a positive feedback loop in which locally translated KIF1C provides additional motor molecules to sustain the persistent and directional transport of its RNA cargoes to locally maintain protrusive extensions during cell movement.

## Supporting information

Supplementary results and Figures

## Acknowledgements

We thank the staff of MRI imaging facility for their technical support. This project was supported by France BioImaging (ANR-10-INBS-04), the Agence Nationale de la Recherche (ANR-11-BSV8-018-02, ANR-14-CE10-0018-01 and ANR-19-CE12-0007-03), and the Ligue Nationale Contre le Cancer and the Fondation pour la Recherche Médicale. This work was supported by the Labex EpiGenMed, from the framework “Investissements d’avenir”. This work was supported in part by the Intramural Research Program of the National Cancer Institute, NIH.

The authors declare no competing financial interests.

## Author contributions statement

EB conceived the study with SM and KZ. Experiments were performed by EB, XP, KM, EC, TiW, MP and RC. AI, KM, SM, ThW and FM analyzed images. EB, XP, KM, SM, EC, ThW, MP, AI, FM and RC analyzed the data. EB, XP, KM prepared the Figures. EB, XP, KM and SM wrote the manuscript.

## Methods

### Generation and maintenance of cell lines

The HeLa-Kyoto cells stably transfected with the KIF1C-GFP BAC were previously described (Maliga et al., 2013; Poser et al., 2008; Chouaib et al., 2020). HeLa Flp-in H9 (a kind gift of S. Emiliani) and the BAC-GFP cells were maintained in Dulbecco’s modified Eagle’s Medium (DMEM, Gibco) supplemented with 10% fetal bovine serum (FBS, Sigma), 100 U/mL penicillin/streptomycin (Sigma) and with 400 μg/ml G418 (Gibco) for the HeLa-Kyoto KIF1C-GFP tagged BAC. NIH/3T3 cells were maintained in DMEM supplemented with 10% calf serum, sodium pyruvate and penicillin/streptomycin at 37°C, 5% CO_2_. Stable cell lines expressing a KIF1C-GFP cDNA were created using the Flp-in system in HeLa Flp-in H9 cells. Flp-in integrants were selected on hygromycin (150 μg ml^−1^). To generate cell lines expressing RNA reporters, NIH/3T3 cells were infected with lentivirus expressing tdMCP-GFP (Addgene plasmid #40649) and GFP-expressing cells with low level of GFP expression were sorted by FACS. This stable population was infected with pInducer20-based reporter constructs expressing β-globin followed by 24xMS2 binding sites and the mouse Net1, Rab13 or control UTRs (pIND20-β24bs/Net1 UTR; pIND20-β24bs/Rab13 UTR; pIND20-β24bs/Ctrl UTR; Moissoglu et al, 2019). Stable lines were selected with geneticin (Thermo Fisher Scientific) and expression of the reporter was induced by addition of 1 μg/ml doxycycline (Fisher Scientific) 2-3 hours before imaging.

For the two-color tracking experiment, NIH/3T3 cells expressing the pIND20-β24bs/Net1 reporter and tdMCP-GFP described below were modified as follows. Stable expression of scFv-mScarletI-GB1 was achieved by lentiviral-mediated integration of a plasmid derived from Addgene (#60906) and sorted by FACS with low level of mScarletI expression. Then a lentivirus expressing a KIF1C fusion with 24 repeats of the GCN4 peptide array was infected in NIH/3T3 cells allowing the detection of KIF1C-ST_x24_ protein with scFv-mScarletI.

### Treatments with siRNAs and drugs

HeLa cells were seeded on 0.17 mm glass coverslips deposited in 6-well plates. Cells were transfected at 70% confluency using JetPrime (Polyplus). Double-stranded siRNAs (30 pmoles) were diluted into 200 μl of JetPrime buffer. JetPrime reagent was added (4 μl) and the mixture was vortexed. After 10 minutes at room temperature (RT), it was added to the cells grown in 2 ml of serum-containing medium. After 24 hours, the transfection medium was replaced with fresh growth medium and cells were fixed 24h later. The sequences of the siRNA were: KIF1C: 5’-CCCAUGCCGUCUUUACCAUdCdG; control: 5’-CAACAGAAGGAGAGCGAAAdTdT. For knockdown of mouse Kif1c the following siRNAs were used, Mm_Kif1c_2 Flexitube siRNA and Mm_Kif1c_3 Flexitube siRNA (Qiagen cat# SI00239687 and SI00239694 respectively) and AllStars negative control siRNA (Qiagen cat# 1027281). siRNAs were delivered into cells using Lipofectamine RNAiMAX (Thermo Fisher Scientific, cat# 13778-150) according to the manufacturer’s instructions. Cells were assayed 72 hours after siRNA transfection.

For drug treatments, 10 μM nocodazole or Cytochalasin D, or an equal volume of DMSO, were added to the growth media for 15 min.

### Plasmid construction

Plasmids were generated with standard molecular biology techniques. Maps and sequences are available upon request.

### Immuno-precipitation and microarrays

HeLa cells containing the KIF1C-GFP BAC were grown to near confluence in 10 cm plates, and two plates were used per IP. Cells were rinsed in ice-cold PBS, and all subsequent manipulations were performed at 4°C. Cells were scraped in HTNG buffer (20 mM HEPES-KOH pH 7.9, 150 mM NaCl, 1% Triton X-100, 10% glycerol, 1 mM MgCl_2_, 1 mM EGTA), containing an antiprotease cocktail (Roche Diagnostic). Cells were incubated for 20 minutes on a rotating wheel, and cellular debris were the removed by centrifugating the extracts 10 minutes at 20,000g. Beads coated with GFP-trap antibody (ChromoTek), or uncoated as control, were washed in HNTG (25 μl of beads per IP). Beads were incubated 1h with a control extract to saturate non-specific binding and then incubated with the proper extract. After 4h of incubation on a rotating wheel, beads were washed four times in HNGT with anti-protease, and twice with PBS. Beads were then incubated with Trizol to extract RNAs, and RNA purification was done as recommended by the manufacturer. The resulting RNAs were amplified and converted into cDNAs by the WT PICO kit (Thermo Fisher), and hybridized on HTA 2.0 chip on an Affymetrix platform (Thermo Fisher). Experiments were performed in duplicates, data were normalized and averaged. Data are deposited on GEO with the following accession number (GSE161316).

### RNA analyses

For total RNA analysis, cells were lysed with Trizol LS reagent (Thermo Fisher Scientific, cat# 10296010) and RNA was extracted according to the manufacturer’s instructions. Isolated RNA was treated with RQ1 RNase-free DNase (M6101, Promega) and analyzed with the nCounter system (NanoString Technologies) using a custom-made codeset. Data were processed using nSolver analysis software (NanoString technologies).

### Single molecule FISH

Cells grown on glass coverslips were fixed for 20 min at RT with 4% paraformaldehyde diluted in PBS, and permeabilized with 70% ethanol overnight at 4°C. For smFISH, we used a set of 44 amino-modified oligonucleotide probes against the GFP-IRES-Neo sequence (sequences given in Table S2). Each oligonucleotide probe contained 4 primary amines that were conjugated to Cy3 using the Mono-Reactive Dye Pack (PA23001, GE Healthcare Life Sciences). To this end, the oligos were precipitated with ethanol and resuspended in water. For labelling, 4 μg of each probe was incubated with 6 μl of Cy3 (1/5 of a vial resuspended in 30 μl of DMSO), and 14 μl of carbonate buffer 0.1 M pH 8.8, overnight at RT and in the dark, after extensive vortexing. The next day, 10 μg of yeast tRNAs were added and the probes were precipitated several times with ethanol until the supernatant lost its pink color. For hybridization, fixed cells were washed with PBS and hybridization buffer (15% formamide in 1xSSC), and then incubated overnight at 37°C in the hybridization buffer also containing 130 ng of the probe set for 100 μl of final volume, 0.34 mg/ml tRNA (Sigma), 2 mM VRC (Sigma), 0.2 mg/ml RNAse-free BSA (Roche Diagnostic), and 10% Dextran sulfate. The next day, the samples were washed twice for 30 minutes in the hybridization buffer at 37°C, and rinsed in PBS. Coverslips were then mounted using Vectashield containing DAPI (Vector laboratories, Inc.).

For smiFISH (Tsanov et al., 2016), 24 to 48 unlabeled primary probes were used (sequences given in Table S2). In addition to hybridizing to their targets, these probes contained a FLAP sequence that was hybridized to a secondary fluorescent oligonucleotide. To this end, 40 pmoles of primary probes were pre-hybridized to 50 pmoles of secondary probe in 10 μl of 100 mM NaCl, 50 mM Tris-HCl, 10 mM MgCl2, pH 7.9. Hybridization was performed at 85°C for 3 min, 65°C for 3 min, and 25°C for 5 min. The final hybridization mixture contained the probe duplexes (2 μl per 100 μl of final volume), with 1X SSC, 0.34 mg/ml tRNA (Sigma), 15% Formamide, 2 mM VRC (Sigma), 0.2 mg/ml RNAse-free BSA, 10% Dextran sulfate. Slides were then processed as above. For Figure S1A, the probes used were RNA and not DNA (sequence in Table S2). The protocol was similar except that hybridization was performed at 48°C and that 50 ng of primary probe (total amount of the pool of probes) and 30 ng of the secondary probes were used per 100 μl of hybridization mix.

For FISH of mouse cells, cells plated on fibronectin-coated coverslips were fixed for 20 min at RT with 4% paraformaldehyde in PBS. FISH was performed with the ViewRNA ISH Cell Assay kit (Thermo Fisher Scientific) according to the manufacturer’s instructions. The flowing probe sets were used: Kif1c (cat# VB6-3200442), Net1 (cat# VB1-3034209), Rab13 (cat# VB1-14374), Ddr2 (cat# VB1-14375), Dynll2 (cat# VB1-18646), Cyb5r3 (cat# VB1-18647). To detect polyA+ RNAs, LNA modified oligodT probes (30 nucleotides) labelled with ATTO 655 were added during hybridization, pre-amplification, amplification and last hybridization steps of the ViewRNA ISH Cell Assay. Cell mask stain (Thermo Fisher Scientific) was used to identify the cell outlines. Samples were mounted with ProLong Gold antifade reagent (Thermo Scientific)

### Imaging of fixed cells

Microscopy slides were imaged on a Zeiss Axioimager Z1 wide-field microscope equipped with a motorized stage, a camera scMOS ZYLA 4.2 MP, using a 63x or 100x objective (Plan Apochromat; 1.4 NA; oil). Images were taken as z-stacks with one plane every 0.3 μm. The microscope was controlled by MetaMorph and figures were constructed using ImageJ, Adobe Photoshop and Illustrator. For the small smiFISH screen, 96-well plates were imaged on an Opera Phenix High-Content Screening System (PerkinElmer), with a 63x water-immersion objective (NA 1.15). Three-dimensional images were acquired, consisting of 35 slices with a spacing of 0.3 μm. FISH images of mouse cells were obtained using a Leica SP8 confocal microscope, equipped with a HC PL APO 63x CS2 objective. Z-stacks through the cell volume were obtained and maximum intensity projections were used for subsequent analysis.

### Image analysis and quantifications

Automated nuclear and cell segmentation was performed with a custom algorithm based on the U-net deep convolutional network (Ronneberger et al., 2015). Nuclear segmentation was performed with the DAPI channel, cell segmentation was performed with the autofluorescence of the actual smFISH image. For segmentation, 3D images were projected into 2D images as described previously (Tsanov et al., 2016). Messenger RNAs were detected with FISH-quant (Mueller et al., 2013) by applying a local maximum detection on LoG filtered images.

For the quantifications of Figure S1D, the distance of every mRNA molecule to the cell membrane was computed as the minimum distance between the mRNA and every point of the cell outline polygon. The distances were then normalized by the square root of the cell area to reduce the impact of the cell size. The plot displays the normalized mean distance of mRNA to the cell membrane for different genes and their standard deviation.

For calculation of Peripheral Distribution Index (PDI) a custom Matlab script was used. The code is described and is available in (Stueland et al., 2019).

### Imaging of live cells

Live imaging (for dual visualization of β24bs/Net1 reporter RNA and KIF1C-ST_x24_ protein) was done using a spinning disk confocal microscope (Nikon Ti with a Yokogawa CSU-X1 head) operated by the Andor iQ3 software. Acquisitions were performed using a 100X objective (CF1 PlanApo λ 1.45 NA oil), and an EMCCD iXon897 camera (Andor). For two-color imaging, samples were sequentially excited at 488 and 540nm. We imaged at a rate of 7.36 fps for 52 sec. The power of illuminating light and the exposure time were set to the lowest values that still allowed visualization of the signal. This minimized bleaching, toxicity and maximized the number of frames that were collected. Cells were maintained in anti-bleaching live cell visualization medium (DMEM^gfp^; Evrogen), supplemented with 10% fetal bovine serum at 37°C in 5% CO_2_ and rutin at a final concentration of 20 mg/l.

Live imaging (for β24bs/Net1 reporter RNA tracking) was done using a Nikon Eclipse Ti2-E inverted microscope, equipped with a motorized stage, a Yokogawa CSU-X1 spinning disk confocal scanner unit, and operated using NIS-elements software. Acquisitions were performed using an Apochromat TIRF 100x oil immersion objective (N.A. 1.49, W.D. 0.12mm, F.O.V. 22mm) and a Photometrics Prime 95B Back-illuminated sCMOS camera with W-view Gemini Image splitter. Constant 37°C temperature and 5% CO_2_ were maintained using a Tokai Hit incubation system. Cells were plated on fibronectin (2mg/ml)-coated 35mm glass bottom dishes and samples were excited using a 488nm (20mw) laser line and imaged at a rate of 6.66 fps for 60 sec.

### Live cell imaging quantification

Images were processed for brightness/contrast, cropped and annotated using ImageJ/FIJI. Kymographs were generated using standard ImageJ/Fiji plugins. Film presentation in figures and videos were edited in ImageJ/Fiji. Bicolor tracking of KIF1C-ST_x24_ proteins and β24bs/Net1 mRNAs was performed using the Manual Tracking plugin in ImageJ/Fiji.

Single color tracking of β24bs/Net1 mRNAs was performed using TrackMate plugin in ImageJ/Fiji. For every cell, all tracks lasting for >2.5 secs (ca. 17 consecutive frames) were used for analysis. Values of ‘Track displacement’ and ‘Linearity of forward progression’ were extracted and plotted. ‘Track displacement’ is defined as the distance from the first to the last spot of the track. ‘Linearity of forward progression’ is the mean straight line speed divided by the mean speed; where mean straight line speed is defined as the net displacement divided by the total track time. For RNA cluster analysis, TrackMate was used to identify spots and extract intensity values. Frequency histograms of spot intensities were plotted using GraphPad Prism software.

## Notes

### Competing Interest Statement

The authors have declared no competing interest.

