## Supplementary results and Figures for "The kinesin KIF1C transports APC-dependent mRNAs to cell protrusions"

### Supplementary results, figures and tables

**Figure S1: Identification of mRNAs transported by the KIF1C motor**

**A**- Localization of mRNAs in protrusion of human cells. Images are micrographs of HeLa cells labelled by smiFISH with probes against endogenous genes. Top: Cy3 fluorescent signals corresponding to endogenous KIF1C, NET1, TRAK2 and RAB13 mRNAs. Bottom: merge panel with red being the Cy3 fluorescent smiFISH signals and blue corresponding to DNA stained with DAPI. Scale bar: 10 microns. Arrowheads indicated accumulation of single molecule mRNA at cell protrusions.

**B**- The KIF1C motor is required for accumulation of RAB13 mRNA in protrusions of HeLa cells. Images are micrographs of HeLa cells labelled by smiFISH with probes against RAB13 mRNAs. Left: Hela cells treated with a control siRNA; right: HeLa cells treated with siRNA against KIF1C. Top: Cy3 fluorescent signal corresponding to endogenous RAB13 mRNA. Lower panels are the merge with Cy3 fluorescent signals in red and DNA in blue (DAPI staining). Scale bar: 10 microns. The orange arrow indicates a protrusion accumulating RAB13 mRNA.

**C**- Efficiency of the KIF1C knock-down measured by smiFISH. The graph depicts the number of single molecules of KIF1C mRNA per HeLa cell, following treatment with control siRNA (orange bar) or KIF1C siRNA (blue bar). Error bars: standard deviation (n=2); number of cells: 50 per condition.

**D**- Quantitative analysis of mRNA localization. The graph depicts the mean distance of mRNAs from the cell membrane, after normalization for cell size. Stars over blue bars: values significantly different than the control mRNAs (KIF20B, PAK2 and MYO18A); stars over orange bars: values significantly different between them. Stars represent p-values: **<0.001, estimated using the Kolmogorov-Smirnov test. Error bars: standard deviation.

**E**- The KIF1C motor preferentially associates with APC-dependent mRNAs. Correlation analysis of mRNA enrichment in protrusions (x axis; Ps/CB is the Protrusion versus Cell Body ratio; data from Wang et al., 2017) versus the enrichment fold of the same mRNAs in KIF1C-GFP IP (y-axis; KIF1C-GFP versus control IP). Red dots depict APC-dependent mRNAs, green dots depict ribosomal protein mRNAs (APC-independent mRNAs), based on data from Wang et al., 2017.

**Figure S2: KIF1C is required for the localization of APC-dependent mRNAs to cytoplasmic protrusions in mouse fibroblasts**

**A**- Micrographs of NIH/3T3 cells labelled by smFISH with probes against the indicated mRNAs. Cells were treated with Kif1C or Control siRNAs. Green outline: cell contour; blue outline: DAPI staining; black spots: mRNAs detected by smFISH. Scale bars: 10 microns.

**B**- Intracellular distribution of *Cyb5r3* and *Kif1c* mRNA calculated by their PDI values, after treatment with Control or Kif1C siRNAs. Horizontal lines represent the mean with 95% confidence interval. Points indicate individual cells. $p$<0.0001, estimated by Student's t-test.

**C**-Localization of APC-independent RNAs is not affected by Kif1c depletion in NIH/3T3 fibroblasts. The graph plots PDI values of *Ddr2*, *Rps20*, *Rpl27a* and polyA+ RNAs, measured from smFISH images of cells treated with Control or Kif1c siRNAs. Points indicate individual cells. Stars are p-values: ****<0.0001, estimated by analysis of variance with Tukey's multiple comparisons test.

**D**- Expression of localized (APC-dependent and -independent) and non-localized mRNAs is not affected by Kif1c depletion in NIH/3T3 fibroblasts. The graph depicts Log2 RNA expression ratios (Kif1c depleted versus control cells), measured by Nanostring analysis. n=3. Bars indicate mean +/- standard error.

**Figure S3: Reporter mRNAs containing Net1 and Rab13 3'UTRs form clusters at the tips of protrusions in mouse fibroblasts**

Micrographs of live NIH/3T3 cells taken 3h after plating and expressing MCP-GFP and the β24bs reporter mRNA carrying the indicated 3' UTRs. Red line: cell contour. Scale bars are 10 microns. Note that reporter mRNAs containing the Net1 or Rab13 3'UTR accumulate in clusters at the tips of protrusions, while the control reporter does not. Boxed insets (right) are magnifications of the areas indicated by the green arrows.

**Figure S4: Reporter mRNAs containing the Net1 3'UTR are transported to protrusions by the by Kif1c motor.**

**A**- Micrograph of a fixed NIH/3T3 cell expressing β24bs/Net1 reporter mRNA, MCP-GFP (green), KIF1C-$ST_{x24}$ protein and scFv-mScarletI (red). Single molecules of β24bs/Net1 mRNAs are visible in green, while single molecules of KIF1C-$ST_{x24}$ protein are red. The numbered white boxes are magnified in B. Blue: DNA stained with DAPI. Scale bar is 5 microns.

**B**- Insets represent magnifications of the boxed areas from the image shown in A. Left: MCP-GFP signals labelling β24bs/Net1 mRNAs; middle: scFv-mScarletI labelling KIF1C-$ST_{x24}$ protein; right: merge with mRNAs in green and KIF1C-$ST_{x24}$ in red. Black and white arrowheads indicate colocalization of single molecules of β24bs/Net1 mRNA and KIF1C-$ST_{x24}$. Scale bars are 5 microns.

**C**- Snapshot of a live NIH/3T3 cell expressing β24bs/Net1 mRNA, MCP-GFP (green), KIF1C-$ST_{x24}$ protein and scFv-mScarletI (red). Snapshot is extracted from Movie 10. The white arrowhead indicates a co-transport event of a single molecule of β24bs/Net1 mRNA (green) with a KIF1C-$ST_{x24}$ protein (Red). The boxed area is zoomed in panels D and E. Scale bar is 5 microns.

**D**- Magnification of the boxed area in panel C, highlighting a co-transport event. Top: KIF1C-$ST_{x24}$; Middle: β24bs/Net1 mRNA; bottom: merged panel with the β24bs/Net1 mRNA in green and KIF1C-$ST_{x24}$ in red. Scale bar is 1 micron.

**E**-Kymograph extracted from Movie 10, showing the trajectory of a single molecule of β24bs/Net1 mRNA (left panel), KIF1C-$ST_{x24}$ (middle panel) and the merge (right panel) in NIH/3T3 cells. Total time of the kymograph is 12.9 seconds and corresponds to the event shown in panels C and D.

**Supplementary spreadsheets:**

**Table S1:** excel file showing the KIF1C-GFP IP microarray data.

**Table S2:** Human smFISH probes sequences and list of probes used for smFISH localization screen in HeLa cells using 26 of the mRNAs most enriched in the KIF1C IP.

**Movie legends:**

**Movie 1:** A live NIH/3T3 cell expressing β24bs/Net1 reporter mRNA and MCP-GFP (green) imaged at 6.67 frames per second for 1 minute, prior to addition of nocodazole. A snapshot of this movie is shown in Figure 3B. Left panel presents the raw signal. Right panel presents the raw signal with overlaid accumulated tracks of individual RNA spots. Only tracks lasting longer than 2.5 seconds are shown. Warmer colors indicate tracks of higher linearity. Scale bar is 5 microns.

**Movie 2:** A live NIH/3T3 cell (same as shown in Movie 1) expressing β24bs/Net1 reporter mRNA and MCP-GFP (green) imaged at 6.67 frames per second for 1 minute, 15 minutes after addition of nocodazole. A snapshot of this movie is shown in Figure 3B. Left panel presents the raw signal. Right panel presents the raw signal with overlaid accumulated tracks of individual RNA spots. Only tracks lasting longer than 2.5 seconds are shown. Warmer colors indicate tracks of higher linearity. Scale bar is 5 microns.

**Movie 3:** A live NIH/3T3 cell expressing β24bs/Net1 reporter mRNA and MCP-GFP (green) imaged at 6.67 frames per second for 1 minute, prior to addition of cytochalasin D. A snapshot of this movie is shown in Figure 3B. Left panel presents the raw signal. Right panel presents the raw signal with overlaid accumulated tracks of individual RNA spots. Only tracks lasting longer than 2.5 seconds are shown. Warmer colors indicate tracks of higher linearity. Scale bar is 5 microns.

**Movie 4:** A live NIH/3T3 cell (same as shown in Movie 3) expressing β24bs/Net1 reporter mRNA and MCP-GFP (green) imaged at 6.67 frames per second for 1 minute, 15 minutes after addition of cytochalasin D. A snapshot of this movie is shown in Figure 3B. Left panel presents the raw signal. Right panel presents the raw signal with overlaid accumulated tracks of individual RNA spots. Only tracks lasting longer than 2.5 seconds are shown. Warmer colors indicate tracks of higher linearity. Scale bar is 5 microns.

**Movie 5:** A live NIH/3T3 cell expressing β24bs/Net1 reporter mRNA and MCP-GFP (green), transfected with control siRNAs, imaged at 6.67 frames per second for 1 minute. A snapshot of this movie is shown in Figure 4A. Left panel presents the raw signal. Right panel presents the raw signal with overlaid accumulated tracks of individual RNA spots. Only tracks lasting longer than 2.5 seconds are shown. Warmer colors indicate tracks of higher linearity. Scale bar is 5 microns.

**Movie 6:** A live NIH/3T3 cell expressing β24bs/Net1 reporter mRNA and MCP-GFP (green), transfected with KIF1C siRNAs, imaged at 6.67 frames per second for 1 minute. A snapshot of this movie is shown in Figure 4A. Left panel presents the raw signal. Right panel presents the raw signal with overlaid accumulated tracks of individual RNA spots. Only tracks lasting longer than 2.5 seconds are shown. Warmer colors indicate tracks of higher linearity. Scale bar is 5 microns.

**Movie 7:** A live NIH/3T3 cell expressing β24bs/Net1 reporter mRNA and MCP-GFP (green), transfected with KIF5B siRNAs, imaged at 6.67 frames per second for 1 minute. A snapshot of this movie is shown in Figure 4A. Left panel presents the raw signal. Right panel presents the raw signal with overlaid accumulated tracks of individual RNA spots. Only tracks lasting longer than 2.5 seconds are shown. Warmer colors indicate tracks of higher linearity. Scale bar is 5 microns.

**Movie 8:** A live NIH/3T3 cell expressing β24bs/Net1 reporter mRNA and MCP-GFP (green), transfected with KIF3A siRNAs, imaged at 6.67 frames per second for 1 minute. A snapshot of this movie is shown in Figure 4A. Left panel presents the raw signal. Right panel presents the raw signal with overlaid accumulated tracks of individual RNA spots. Only tracks lasting longer than 2.5 seconds are shown. Warmer colors indicate tracks of higher linearity. Scale bar is 5 microns.

**Movie 9:** A live NIH/3T3 cell expressing β24bs/Net1 reporter mRNA, MCP-GFP (green), KIF1C-ST$_{x24}$ protein and scFv-mScarletI (red) was imaged at 7.36 frames per second during 6.45 seconds. The movie follows a co-directed transport event highlighted by the white arrowhead in Figure 6C-E. Scale bar is 5 microns. Top left insets are magnifications of the yellow boxed area showing separately each fluorescent channel. Scale bar is 1 micron.

**Movie 10:** A live NIH/3T3 cell expressing β24bs/Net1 reporter mRNA, MCP-GFP (green), KIF1C-ST$_{x24}$ protein and scFv-mScarletI (red) was imaged at 7.36 frames per second during 12.9 seconds. The movie follows a co-directed transport event highlighted by the white arrowhead in Figure S4C-E. Scale bar is 5 microns. Top left insets are magnifications of the yellow boxed area showing separately each fluorescent channel. Scale bar is 1 micron.

Fig. S1

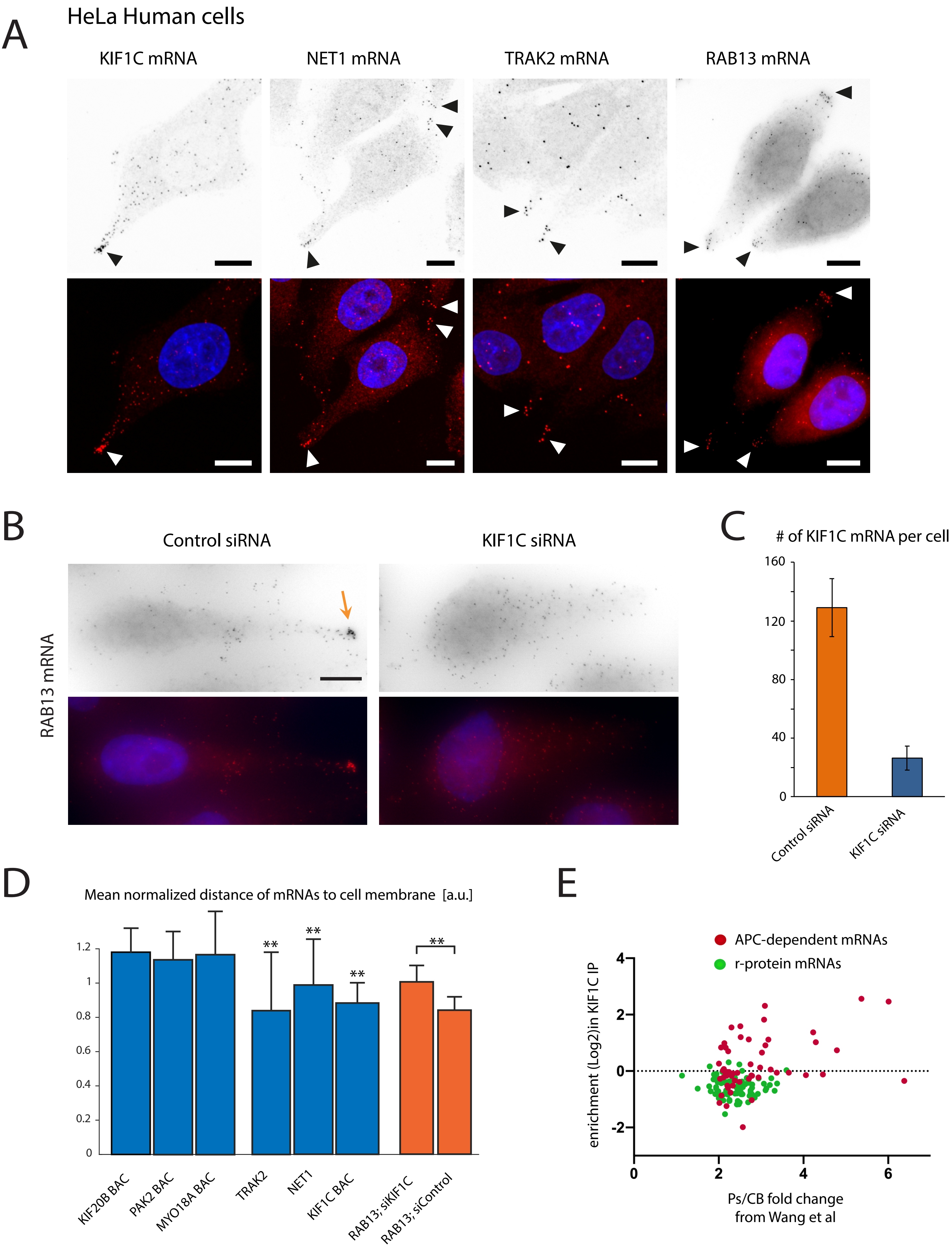
A
HeLa Human cells
KIF1C mRNA
NET1 mRNA
TRAK2 mRNA
RAB13 mRNA
B
Control siRNA
KIF1C siRNA
RAB13 mRNA
C
### of KIF1C mRNA per cell
160
120
80
40
0
Control siRNA
KIF1C siRNA
D
Mean normalized distance of mRNAs to cell membrane [a.u.]
1.2
1
0.8
0.6
0.4
0.2
0
**
**
**
**
KIF20B BAC
PAK2 BAC
MYO18A BAC
TRAK2
NET1
KIF1C BAC
RAB13; siKIF1C
RAB13; siControl
E
APC-dependent mRNAs
r-protein mRNAs
enrichment (Log2)in KIF1C IP
4
2
0
-2
0
2
4
6
Ps/CB fold change
from Wang et al

Fig. S2

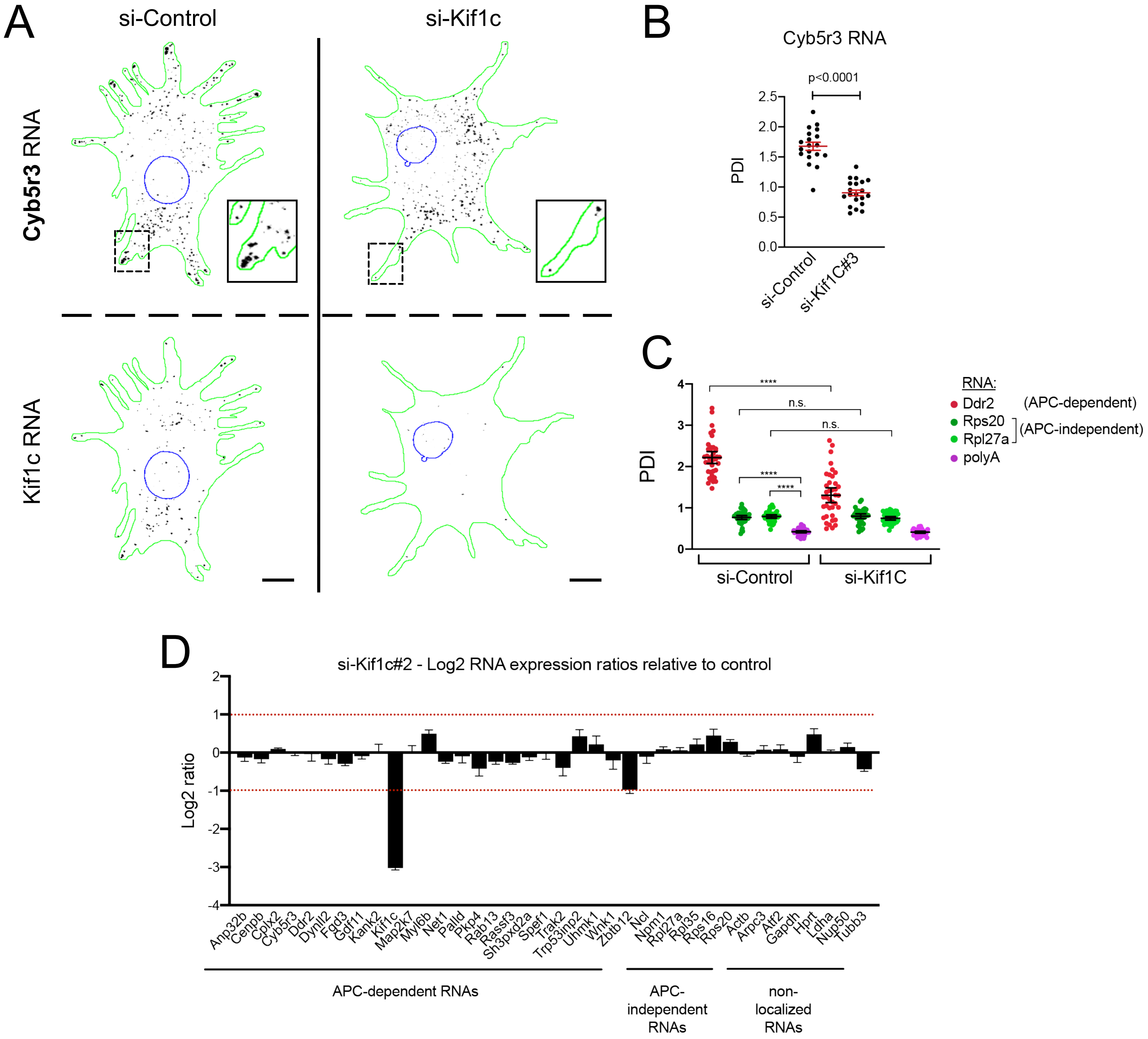
A
si-Control
si-Kif1c
Cyb5r3 RNA
Kif1c RNA
B
Cyb5r3 RNA
p<0.0001
PDI
2.5
2.0
1.5
1.0
0.5
0.0
si-Control
si-Kif1C#3
C
PDI
4
3
2
1
0
****
n.s.
n.s.
****
****
si-Control
si-Kif1C
RNA:
Ddr2 (APC-dependent)
Rps20
Rpl27a
(APC-independent)
polyA
D
si-Kif1c#2 - Log2 RNA expression ratios relative to control
Log2 ratio
2
1
0
-1
-2
-3
-4
Anp32b
Cenpb
Cplx2
Cyb5r3
Ddr2
Dynll2
Fgd3
Gdf11
Kank2
Kif1c
Map2k7
Myl6b
Net1
Palld
Pkp4
Rab13
Rassf3
Sh3pxd2a
Spef1
Trak2
Trp53inp2
Uhmk1
Wnk1
Zbtb12
Ncl
Npm1
Rpl27a
Rpl35
Rps16
Rps20
Actb
Arpc3
Atf2
Gapdh
Hprt
Ldha
Nup50
Tubb3
APC-dependent RNAs
APC-independent RNAs
non-localized RNAs

Fig. S3

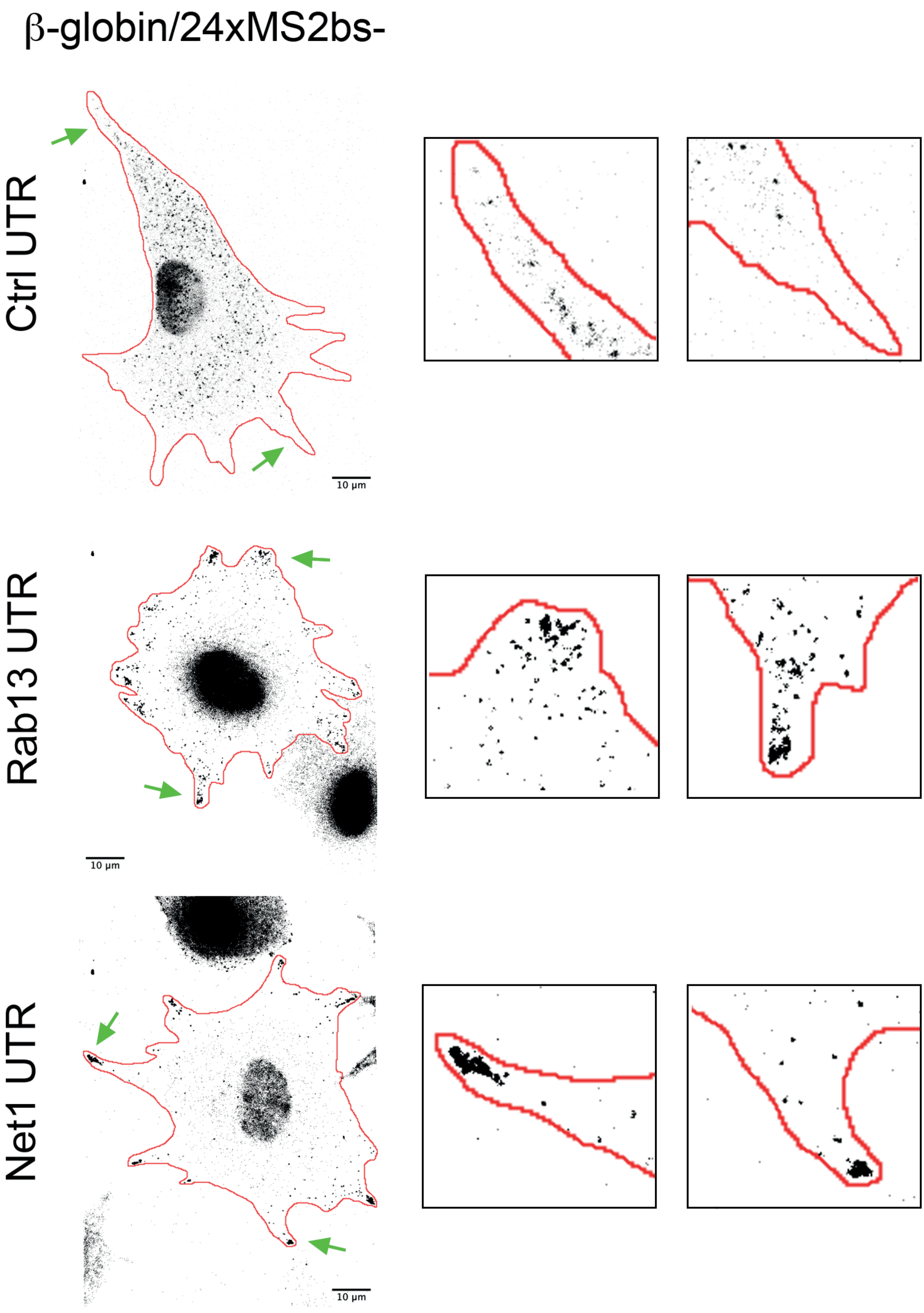

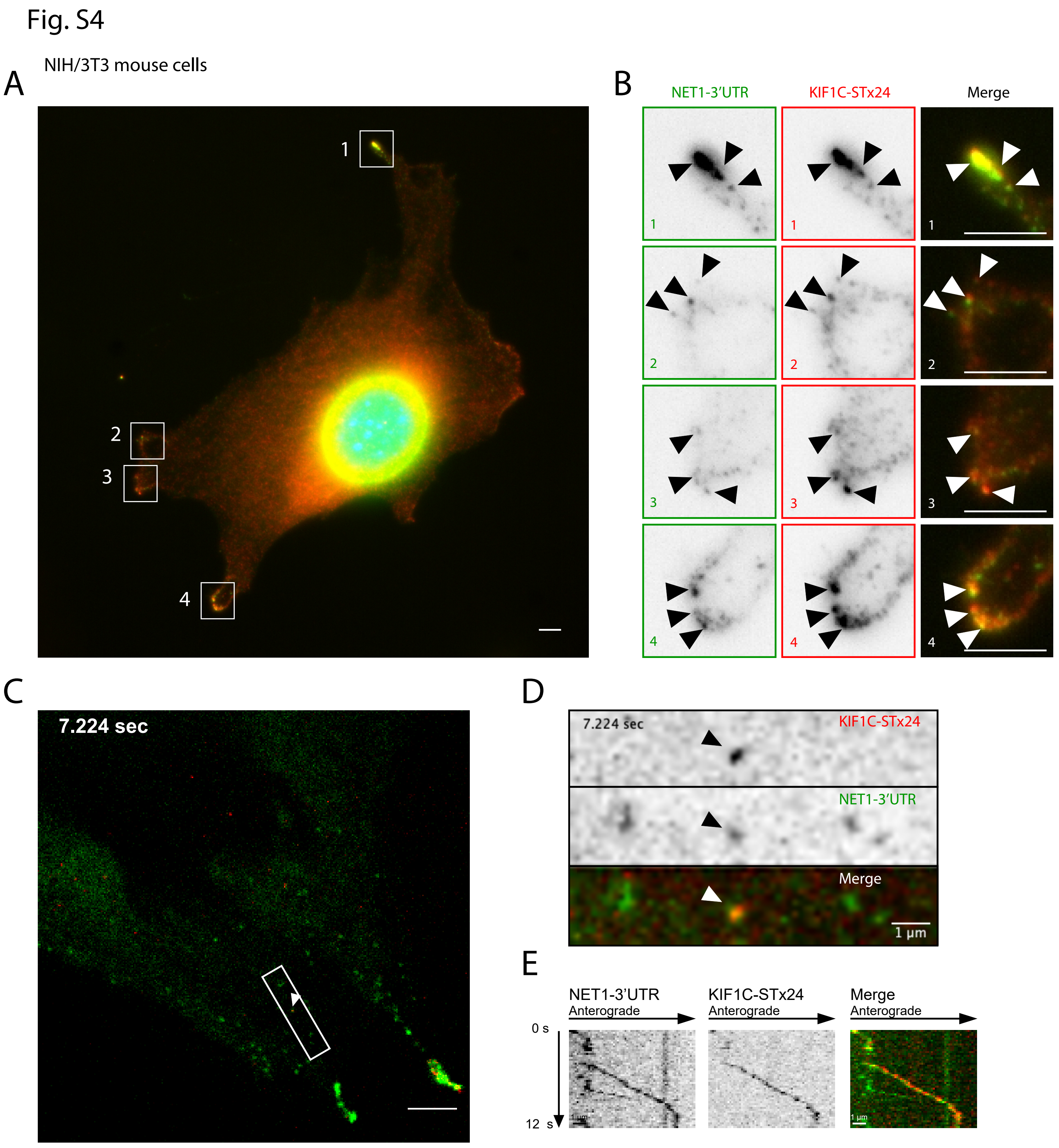
Fig. S4
A
NIH/3T3 mouse cells
1
2
3
4
B
NET1-3'UTR
KIF1C-STx24
Merge
C
7.224 sec
D
7.224 sec
KIF1C-STx24
NET1-3'UTR
Merge
1 µm
E
NET1-3'UTR
Anterograde
KIF1C-STx24
Anterograde
Merge
Anterograde
0 s
12 s
